# Modelling the role of contour integration in visual inference

**DOI:** 10.1101/2022.10.28.514169

**Authors:** Salman Khan, Alexander Wong, Bryan Tripp

## Abstract

Under difficult viewing conditions, the brain’s visual system uses a variety of recurrent modulatory mechanisms to augment feed-forward processing. One resulting phenomenon is contour integration, which occurs in the primary visual (V1) cortex and strengthens neural responses to edges if they belong to a larger smooth contour. Computational models have contributed to an understanding of the circuit mechanisms of contour integration, but less is known about its role in visual perception. To address this gap, we embedded a biologically grounded model of contour integration in a task-driven artificial neural network, and trained it using a gradient-descent variant. We used this model to explore how brain-like contour integration may be optimized for high-level visual objectives as well as its potential roles in perception. When the model was trained to detect contours in a background of random edges, a task commonly used to examine contour integration in the brain, it closely mirrored the brain in terms of behavior, neural responses, and lateral connection patterns. When trained on natural images, the model enhanced weaker contours and distinguished whether two points lay on the same vs. different contours. The model learnt robust features that generalized well to out-of-training-distribution stimuli. Surprisingly, and in contrast with the synthetic task, a parameter-matched control network without recurrence performed the same or better than the model on the natural-image tasks. Thus a contour integration mechanism is not essential to perform these more naturalistic contour-related tasks. Finally, the best performance in all tasks was achieved by a modified contour integration model that did not distinguish between excitatory and inhibitory neurons.

**Author summary:** Deep networks are machine-learning systems that consist of interconnected neuron-like elements. More than other kinds of artificial system, they rival human information processing in a variety of tasks. These structural and functional parallels have raised interest in using deep networks as simplified models of the brain, to better understand of brain function. For example, incorporating additional biological phenomena into deep networks may help to clarify how they affect brain function. In this direction, we adapted a deep network to incorporate a model of visual contour integration, a process in the brain that makes contours appear more visually prominent. We found that suitable training led this model to behave much like the corresponding brain circuits. We then investigated potential roles of the contour integration mechanism in processing of natural images, an important question that has been difficult to answer. The results were not straightforward. For example, the contour integration mechanism actually impaired the network’s ability to tell whether two points lay on the same contour or not, but improved the network’s ability to generalize this skill to a different group of images. Overall, this approach has raised more sophisticated questions about the role of contour integration in natural vision.

## Introduction

Deep neural networks (DNN) are often used as models of the visual system [1–6]. It has been argued that they are mechanistic models [6] because some of their computational elements have analogies in the brain. But they lack many other biological mechanisms, which may contribute to differences in representations [5, 7, 8] and behavior [9–15]. In contrast, there are many physiological models of circuits that underlie localized neural phenomena, such as [16–22], but these models tend to be isolated from larger circuits and to have uncertain connections with ethologically important visual tasks.

The limitations of both deep networks and isolated circuit models might potentially be addressed by combining them, i.e. incorporating detailed circuit models into deep networks. In this direction, recent studies have incorporated details of interlaminar and interareal connectivity into deep networks [8, 23–25]. Few studies [17, 26–29] have incorporated biologically grounded microcircuits into functionally sophisticated deep networks, but doing so may be an important step in understanding how microcircuits contribute to behaviour, and reproducing the superior generalization abilities of the brain [30].

Contour integration [31–34] is a phenomenon in the V1 cortex where stimuli from outside a neuron’s classical receptive field (cRF) modulate its feed-forward responses (Fig. 1). In particular, a neuron’s response is enhanced if a preferred stimulus within the cRF is part of a larger contour. Li et al. [32] found that these elevated V1 responses were highly correlated with contour detectability. Under difficult viewing conditions, it is thought that the that the visual system uses contour integration to *pop out* smooth contours. Contour integration is mediated by intra-area lateral and higher-layer feedback connections [35, 36]. Past computational models [22, 37–40] have tested potential mechanisms and successfully replicated neurophysiological data. However, a limitation of all of these circuit models is that they are stand-alone models that do little to clarify the roles of contour integration in natural vision.

**Fig 1.**
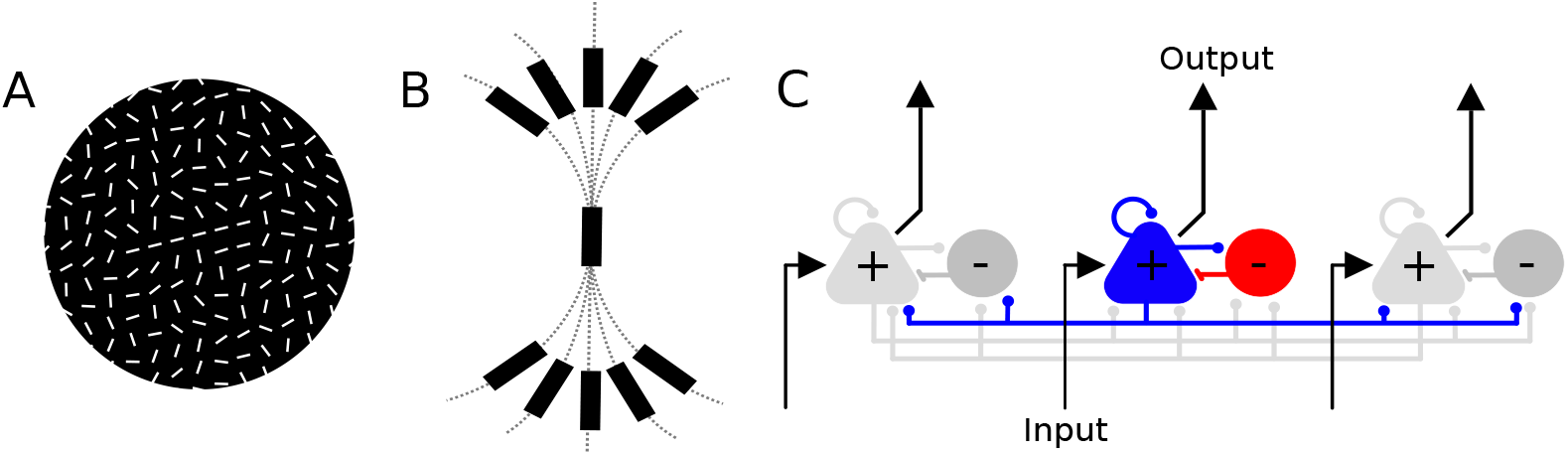
Contour integration. A: Contour integration has been studied using stimuli in which line segments form a contour within a larger field of randomly oriented segments. Subjectively, the contour pops out from the background. B: Contour integration is thought to be mediated by long range inter-area and feedback connections from higher layers. The Association Field Model [31] is commonly used to model intra-area lateral connections. These long-range connections preferentially connect neurons with co-linear or co-circular orientation preferences. C: Microcircuit architecture of the circuit model [22] on which our work is based, which focuses on the role of lateral connections in V1. The outgoing connections of one of the excitatory nodes are highlighted in blue, while those of its paired inhibitory node are shown in red. Connections ending in a circle are excitatory while those ending in a bar are inhibitory.

In this work, we embedded a circuit model of contour integration within a deep network. We used this model to investigate two broad questions. First, we tested whether key characteristics of biological contour integration would emerge as a result of the network learning to identify contours within backgrounds of randomly oriented edges (a kind of stimulus that has often been used to study contour integration). We found that the trained model was consistent with biological data on behaviour (detection of contours), electrophysiology (unit responses versus contour length and contour-fragment spacing), and connectivity (structure of learnt lateral connections). This provides new evidence that these particular circuit characteristics benefit the perception of contours within these synthetic visual stimuli. Second, we used our model to investigate whether contour integration improved performance of two natural-scene tasks. One of these was detection of weak edges in natural scenes, a role that has previously been proposed for contour integration. The second was a new task that required distinguishing connected contours from nearby unconnected contours. In the first task, the contour integration model performed similarly to a parameter-matched feed-forward network. In the second task, surprisingly, the model performed much worse than the control network. However, it generalized better to a variation of the task that it was not trained on. Furthermore, a variation of the model that allowed excitatory neurons to inhibit some of their targets substantially outperformed the control. This suggests that contour integration is relevant to the second task, but the model we adopted was not optimal, either because biological contour integration is not optimal or because important biological elements were missing from the model.

## Model

### Contour integration block

We adapted an existing circuit model of V1 contour integration and incorporated it into an artificial neural network (ANN). We used the current-based-subtractive-inhibition model of Piech et al. [22]. This model focuses on within-layer lateral interactions between V1 orientation columns (co-located populations of neurons that respond to edges of similar orientations over a small area of visual space). Piech et al. modelled each orientation column using a pair of reciprocally connected excitatory (E) and inhibitory (I) nodes whose temporal dynamics were defined as,

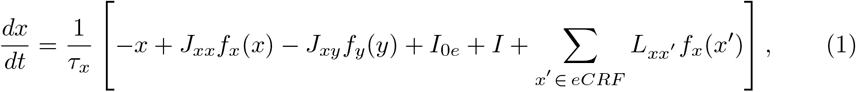

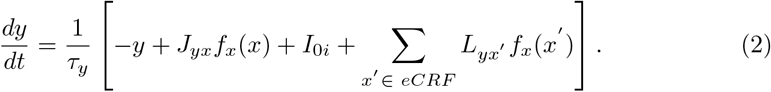

Here, *x* and *y* are membrane potentials of E and I nodes, *J_xx_, J_xy_, J_yx_* are E→E (self), I→E and E→I within node-pair connection strengths, respectively, *f*.(.) is a non-linear activation function that transforms membrane potentials into firing rates, *τ*. are membrane time constants, *I* is the external input current to the model, *I*_0_. are background currents and *L_xx′_* and *L_yx′_* are strengths of lateral connections from E nodes *x′* in neighboring columns that lie within the extra classical receptive field (e-cRF).

An E-I pair of this model and all of its connections are shown in Fig 1C. E nodes process incoming edge extraction responses while I nodes subtractively modulate E node activities. Nodes also receive recurrent inputs from nearby columns via lateral connections. Piech et al. designed these anisotropically distributed connections [41] with connectivity patterns suggested by the Association Field Model [31] (Fig. 1B); each column has the most dense lateral excitatory connections with nearby columns sensitive to edge fragments that are co-linear with the column’s preferred orientation. A similar but orthogonally oriented association field was used to model inhibitory connections.

Piech et al. defined the full model over a 2D grid of spatial locations. Each spatial location contained a set of orientation columns with the same frequency selectivities and a range of orientation preferences. The lateral connections of each orientation column were hard-coded. The dynamics of the full model were realized as the joint activities of all columns.

We made minimal adaptations to this circuit model to implement it as a trainable block inside a convolutional network. First, we replaced summations over e-cRFs with convolutions. The convolution operates over columns in nearby locations as well as at the same location. It incorporates the excitatory self connection, *J_xx_*, and the lateral connections. Second, we used Euler’s method to express the dynamics as difference equations [28, 42]. Third, we defined all model parameters including the lateral connections to be learnable and used task-level optimization to learn their optimal settings. Piech et al. [22] distinguished excitatory and inhibitory neurons in their model, consistent with Dale’s principle [43, 44]. In contrast, neurons in convolutional networks typically do not make this distinction, but allow weights to take on whatever values maximize performance. This consistently results in each neuron exciting some of its targets and inhibiting others. To ensure individual nodes were consistent with Dale’s principle, we constrained weights to be positive or negative, as appropriate. For connections between paired excitatory and inhibitory neurons, a logistic sigmoid non-linearity was applied to the learned weight parameter to prevent changes in sign. The same method was used to retain the sign of the model’s time constants. For lateral connection kernels, a positive-only constraint was imposed on each element during training.

With these modifications, the activity of an orientation column is expressed as,

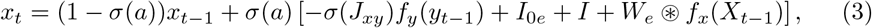

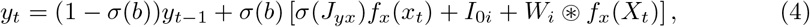

where *x*_0_ = *y*_0_ = 0.

Here, *x* and *y* are membrane potentials, *f*.(.) is a non-linear activation function, *σ*(*a*), *σ*(*b*) are membrane time constants, *σ*(*J_xy_*), *σ*(*J_yx_*) are local I → E, E → I connection strengths, *σ*() is the logistic sigmoid function which constrains time constants and local connection strengths to be positive, *W_e_* are lateral excitatory connections from E nodes in nearby columns to E, *W_i_* are connections from nearby E nodes to I, *f_x_*(*X_t_*) is the output of all modeled nodes at time *t*, ⊛ is the convolution operator, *I* is the external input and *I*_0_. is a node’s background activity.

This final form is a recurrent neural network that can be trained using standard neural network training techniques [42]. Lastly, we included a batch normalization (BN) [45] layers after every convolutional layer to model weak omni-directional inhibition [46]. We refer to this transformed model as the contour integration (CI) block and include it as a whole inside ANNs. Parameters of the CI block and their settings are described in the Methods Section.

### Visual inference network

The full model is composed of edge extraction, CI and classification blocks (see Fig. 2). For edge extraction, we used the first convolutional layer of a ResNet50 [47] that was pre-trained on the ImageNet [48] dataset. We additionally added BN and max-pooling after the convolutional layer in all tasks other than edge detection in natural images. This helped reduce computational complexity (by reducing the spatial dimensions over which the recurrent CI block acts) and improved performance as well. For the task of edge detection in natural images, only the BN was added. Outputs of the edge extraction block were fed into the CI block. The same CI block was used across all tasks.

**Fig 2.**
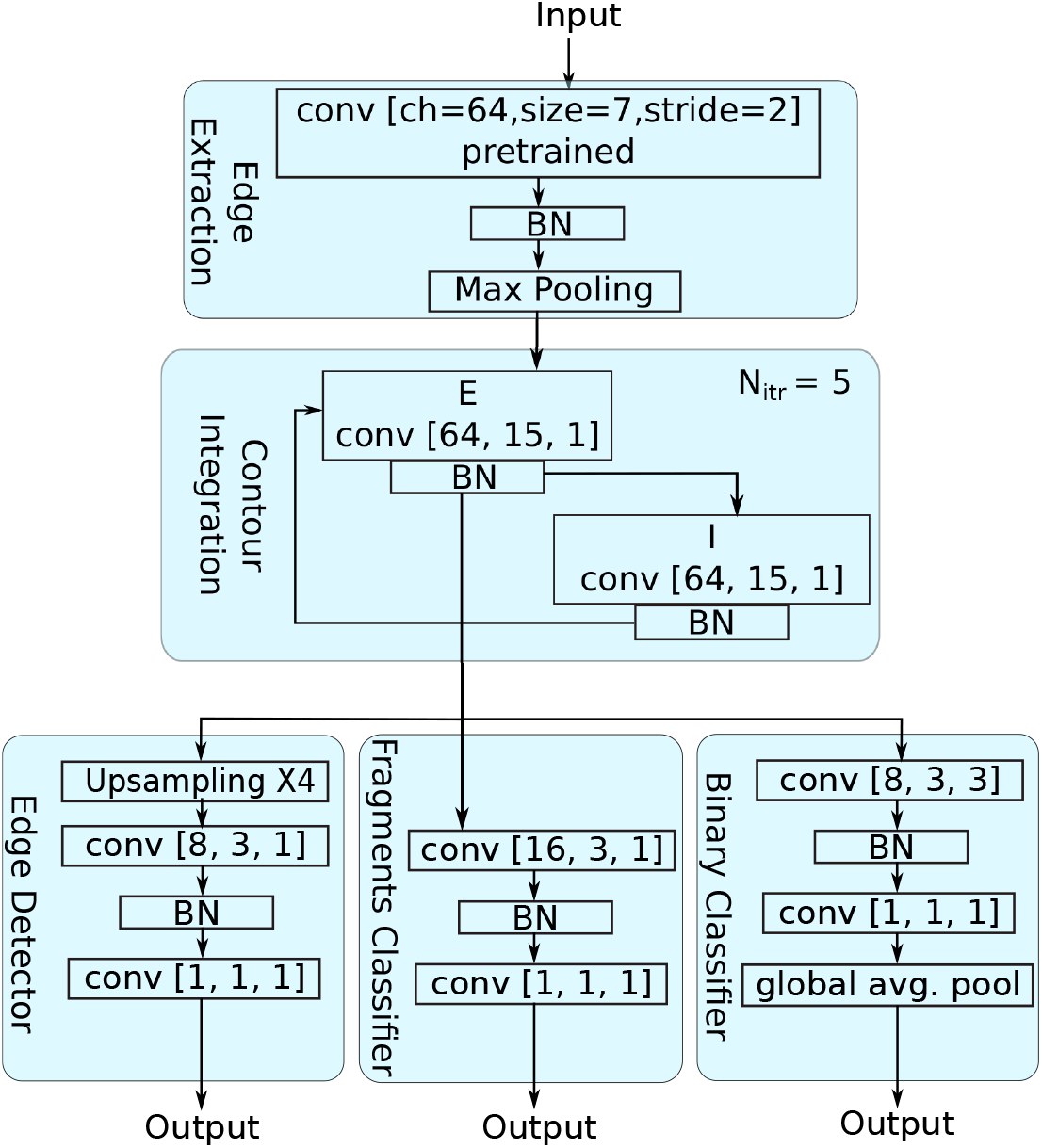
Model Architectures. The main component of the model is the contour integration (CI) block. It consists of a 3D grid of orientation columns and models the horizontal interactions between them. Each orientation column is modeled by a pair of excitatory (E) and inhibitory (I) nodes. Each orientation column receives as input the output of an edge extraction unit at the same spatial location and channel. Horizontal connections connect orientation columns with other orientation columns at different spatial locations and channels. These connections are learnt by optimizing performance on high-level tasks. The full model consists of three main blocks: edge extraction, CI and classification blocks. Edge extraction and CI blocks are common for all tasks. For edge extraction, the first convolutional layer of a ResNet50 [47] that was previously trained on ImageNet [48] was used. Task specific classification blocks (edge detector, fragments classifier, binary classifier) map contour integration activations to output labels. For each convolutional (conv) layer, the square brackets specify the number, size, and stride length of kernels. Batch normalization (BN) layers were typically used after convolutional blocks. Bi-linear interpolation was used for up-sampling in the edge detector classification block.

Outputs of the CI blocks were passed to classification blocks. Classification blocks mapped CI block outputs to required label sizes for each tasks. These blocks had two convolutional layers each. Deeper classification blocks might have allowed better task performance, but we chose shallower classification blocks so that the CI block would play an essential role in network function. Description of each of the classification blocks can be found in the Methods Section. The architectures of the all the models we used are shown in Fig. 2.

#### Feedforward control network

We compared our contour-integration model (the visual inference network described above) with a feed-forward control network of matching capacity (number of parameters). Feed-forward networks can be parameterized to match capacity in several different ways [49, 50]. Because we were interested in modeling V1 lateral connections, we used convolutional kernels of the same size as the model. Compared to standard convolutional kernels, these were much larger and were specifically designed to model lateral connections, which may spread out up to eight times the cRF of V1 neurons [41].

The control network used the same edge extraction and classification blocks as the model. Only the middle block was different. The control’s middle block used the same convolutional layers as the model’s CI block but ordered them sequentially. Additionally, batch normalization and dropout layers (*p_dropout_* = 0.3) were added after every convolutional layer to prevent the control from over-fitting the training data. Finally, no positive-only weight constraint was enforced on the control network. It was free to adopt any weight changes that improved performance. Compared to the control, the CI block does ≈ *N_iter_* more computations per image and has a larger inference run time because it is recurrent. However, this is consistent with contour integration in the brain, which affects late-phase responses of V1 neurons rather than their initial responses [32].

## Results

### Contour detection

We first trained the networks with stimuli that are typically used to study biological contour integration [31, 32, 51]. These stimuli consist of many small edges, a few aligned to form a contour, and the rest randomly oriented to form the background (see Fig. 3). Li et al. [51] found that macaque monkeys progressively improved at detecting contours and had higher contour-enhanced V1 responses with experience on these stimuli. Hence, contour integration is learnable from these stimuli.

**Fig 3.**
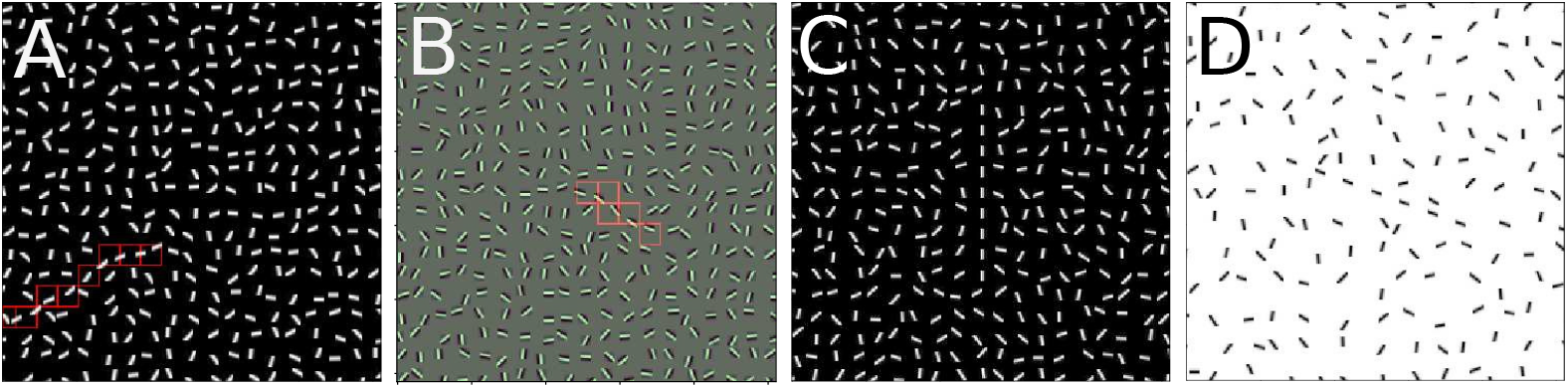
Contour fragments stimuli. A and B, Example training stimuli. All line segments are identical Gabor fragments. A few adjacent fragments were aligned to form a smooth contour (highlighted in red). Remaining fragments were randomly distributed. Embedded contours differed in their location, length, inter-fragment curvature and their component Gabors. C and D, Test stimuli use to analyze the impact of length and inter-fragment spacing. Test stimuli consisted of centrally located straight contours with different lengths (C) and different spacing between contour fragments (D).

We constructed a dataset containing 64,000 training and 6,400 validation images in which contours differed in their locations, lengths *l_c_* (number of edge fragments that made up the contour), inter-fragment degree of curvature *β*, and edges (Gabor functions with different parameters). Details of the full dataset are described in the Methods Section.

Networks were tasked with identifying fragments that were part of the contour. A fragments classifier block (see Fig. 2) followed the CI block to map its outputs to the desired label size. Details of the training process are described in the Methods Section. Network performances were evaluated using mean Intersection over Union (IoU) scores between predictions and labels (see Methods Section). We refer to this task as contour detection due to its similarity with object detection in computer vision, but note that it differs from the kind of detection used in monkey experiments, which involves two patches of line segments and requires only selection of the patch that contains a contour [51].

Averaged peak IoU scores over training are shown in Table 1. For each network, results were averaged over five independent runs that were initialized with different random seeds. The model outperformed the control by ≈ 11% (validation score).

**Table 1.** Peak IoU scores on the contour fragments dataset. Peak values (mean ± 1 SD) were averaged across five independent runs for each network.

| Network | Train (%) | Validation (%) |
| --- | --- | --- |
| Model | $87.33 \pm 0.28\%$ | $84.48 \pm 0.30\%$ |
| Control | $71.62 \pm 0.35\%$ | $73.61 \pm 0.38\%$ |

#### Effect of contour length and inter-fragment spacing

To determine whether networks learnt to integrate contours in a manner similar to the brain, we analyzed them for consistency with behavioral and neurophysiological data. Li et. al. [32] concurrently monitored behavioral performance and V1 neural responses of macaque monkeys as the length of embedded contours and the spacing between contour fragments were varied. At the behavioral level, contours became more salient as lengths increased. Furthermore, when contours extended in the direction of the preferred orientation of V1 neurons, firing rates monotonically increased. Conversely, when spacing between fragments increased, contours became less salient and V1 firing rates decreased monotonically.

In a similar manner, we analyzed trained networks behaviorally at the output of networks and neurophysiologically at the output of centrally located neurons of the CI blocks. For the contour-integration model network, this corresponded to the outputs of E neurons while for the control network it corresponded to the outputs of the second convolutional layer. Behavioural performance was quantified using task-level mean IoU scores. Similar to [32], neurophysiological responses were quantified by the contour integration gain,

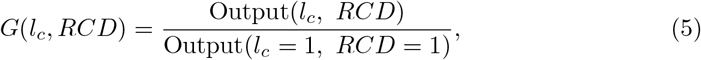

where the relative co-linear distance (RCD) quantifies inter-fragment spacing and was defined as the ratio of inter-fragment spacing to fragment length in pixels. The condition *l_c_* = 1, RCD=1 is when a neuron receives its optimal stimulus within its cRF and no neighboring contour fragments align with it.

We constructed separate test stimuli (similar to those of [32]) for each recorded neuron. These consisted of centrally located contours of varying length and inter-fragment spacing, where each contour fragment was a spatially shifted copy of the neuron’s optimal within-cRF stimulus. A detailed description of the test stimuli is given in the Methods Section. Examples are shown in Fig. 3C and D.

Average IoU scores as contour length increased are shown in Fig. 4A. Results were averaged over five copies of each network, each trained in the same way but initialized with different random weights. For centrally located straight contours, behavioural performance of both networks was similar. Both the contour-integration model and control networks excelled (≥ 95%) at detecting the absence of contours. There were dips in performance for length-three contours as they were the hardest to detect. For all other lengths, prediction accuracy increased with length with the model outperforming the control at larger contour lengths.

**Fig 4.**
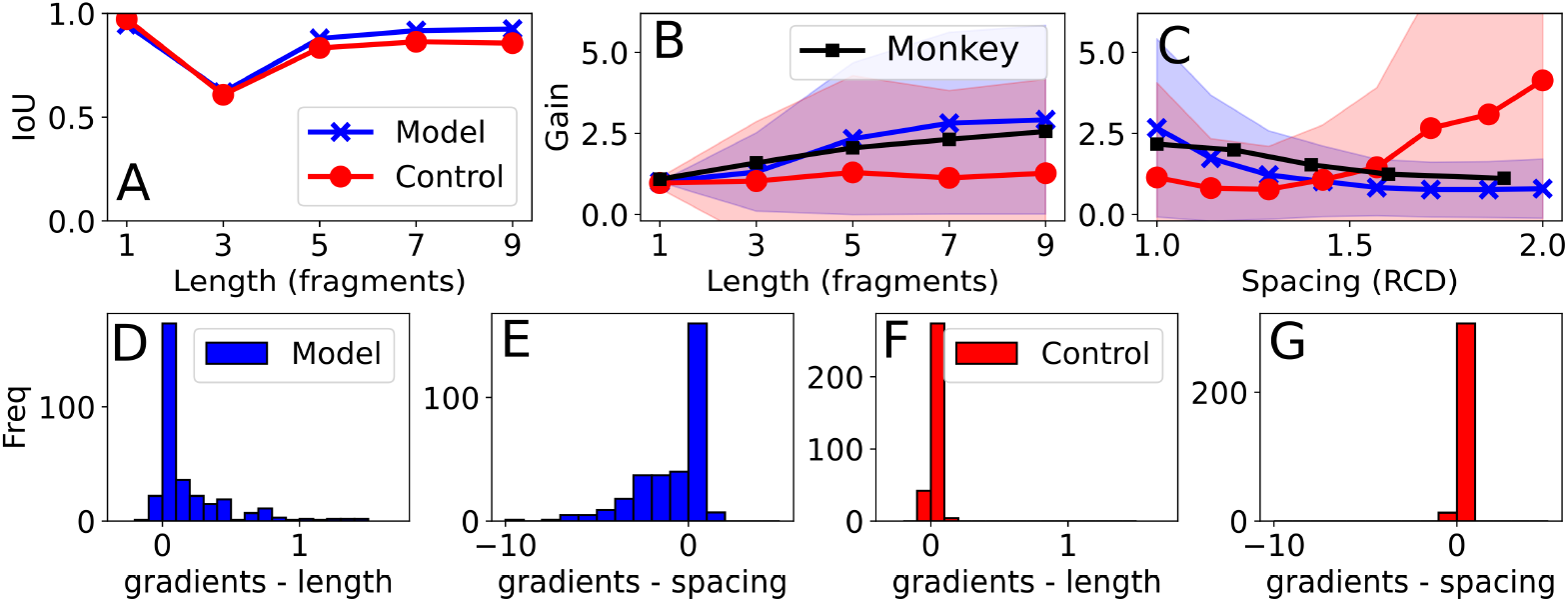
Synthetic contour fragments results. A, IoU vs. contour length for straight contours. Behavioural classification performance of the model and control were similar. B and C, Population average gains vs. length and vs. fragment spacing, respectively. Contour lengths are expressed in number of fragments and spacing between fragments are expressed as relative co-linear distance (RCD). RCD is defined as the ratio between the spacing between fragments to the length of a fragment. Neurophysiological results are from Li et al. [32]. The plot shows the weighted average gains from the two monkeys used in their study. Dark lines show mean values and shaded areas represent unit standard deviation from means, over neurons from five different training runs for each model. D and E, Gradients of linear fits of the outputs of individual neurons as contour length and as inter-fragment spacing were increased. F and G, similar plots as D and E but for the control. The contour-integration model showed consistent trends with neurophysiological data while the control behaved differently.

Larger contrasts between the model and control were observed when neural response gains were analyzed. Fig. 4B shows population average gains as contour lengths changed, along with averaged gains from two monkeys in Li et al. [32]. In the contour-integration model network, average gains increased monotonically with contour length, similar to the monkey data. In contrast, average gains in the control network did not change appreciably with contour length. Fig. 4C shows population average gains as the spacing between fragments increased. Model-network gains decreased monotonically with spacing, consistent with the monkey data [32]. Control-network gains, unexpectedly, increased with spacing.

To calculate gains in both the model and the control network, we excluded neurons that did not respond to any single Gabor fragment in the cRF (no optimal stimulus). Out of the 320 possible neurons, 188 model and 178 control neurons were retained according to this criterion. Furthermore, for population average gains, neurons that were unresponsive to any contour condition (all zero gains) and those that had outlier gains (≥ 20) for any contour condition were also removed. Typically, these large gains were seen for neurons that had small responses to *l_c_* = 1 contours and small changes in the CI block outputs significantly affected their gains. This resulted in the removal of an additional 36 model and 144 control neurons. Across each population (model and control), there was a wide range of enhancement gains exhibited by individual neurons as shown in the mean ±1 SD shaded area in Fig. 4B.

To better understand how responses varied across neuron populations, we plotted histograms of the slopes of linear fits to CI block outputs versus contour length and inter-fragment spacing. This was done for all neurons for which the optimal stimulus was found. Since outputs rather than gains were considered, we included neurons with outlier gains in these histograms. Results of the model network are shown in Fig. 4D and E while those of the control network are shown in Fig. 4F and G. Most model neurons showed positive slopes as contour lengths increased and negative slopes as fragment spacing increased, consistent with trends in the monkey data. In contrast, the slopes of control-network responses versus fragment length and spacing were both clustered slightly above zero. While the task performance of the model and the control networks were similar, they employed different strategies to solve the task, and only the contour-integration model network was consistent with neurophysiological data.

#### Lateral connectivity patterns

We also analyzed learnt lateral kernels for consistency with neuroanatomical properties of V1 lateral connections. The model constrained the sign of lateral kernels to be positive, consistent with Dale’s principle and the model of [22]. We used separate kernels to model excitatory connections onto excitatory neurons and inhibitory neurons. As in [22], inhibitory neurons only synapsed onto neurons in the same column. Examples of learnt lateral kernels are shown in Fig.5A and B. A full set of learnt lateral kernels of a trained model is shown in S1 Fig, S2 Fig and S3 Fig. Qualitatively, most excitatory-targeting connections were anisotropically distributed and spread out densely in the preferred orientations of the source neurons, while inhibitory-targeting connections were shorter and more omni-directional.

**Fig 5.**
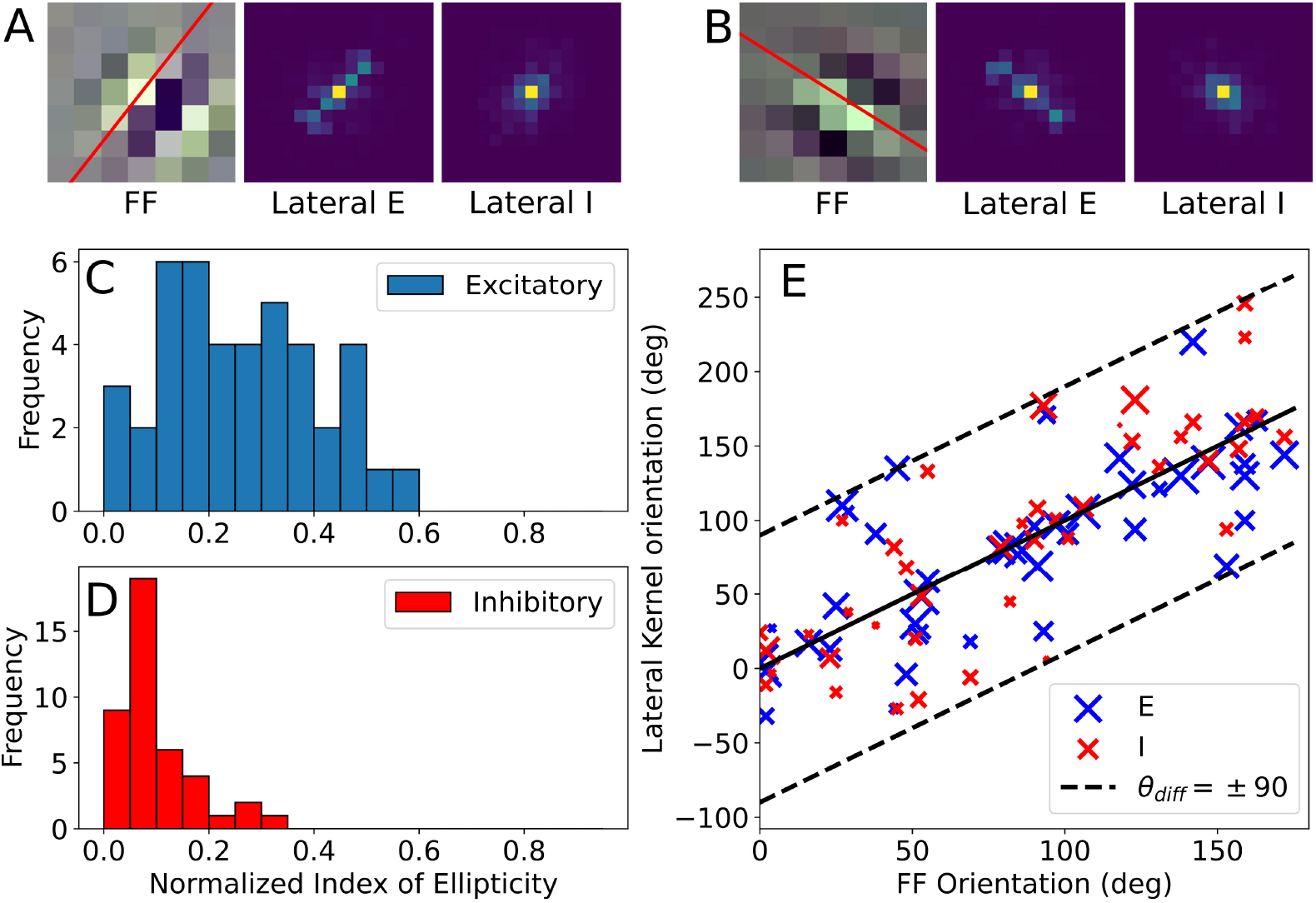
lateral kernel analysis results. A and B, Example learnt excitatory (E) and inhibitory (I) lateral connections. The leftmost subplot shows a kernel in the feedforward (FF) edge extraction layer. The red line through its center shows its preferred orientation. The middle subplot shows its corresponding learnt lateral E connections while the rightmost subplot shows its learnt lateral I connections. Each lateral kernel had 64 channels. To visualize the kernels, the channel dimension was summed out. C and D, Histograms of normalized index of ellipticity of lateral E and I connections respectively. Lateral E connections spread out further and are more directed than inhibitory connections. C, Axis-of-elongation of lateral connections plotted against the orientation of their corresponding feedforward edge extraction kernels. Each point is scaled by its normalized index of ellipticity; larger markers are more directed kernels. Dashed lines show ±90° angular difference. Lateral kernels that lie on these lines are orthogonal to feedforward kernels.

We quantified the spread of lateral connections using a procedure adapted from [52]. Sincich and Blasdel injected axon staining dye into V1 orientation columns, and characterized the staining pattern around each injection with an averaging vector, 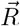. The magnitude, *r*, indicated the direction selectivity of lateral connections, while its angle pointed in the direction of the densest staining. They also calculated a normalized index of ellipticity *r_n_* by normalizing r with the mean length of all lateral connections vectors (see Methods Section for details). A *r_n_* of zero indicates an omni-directional spread of lateral connections while a value of one indicates a straight line. Finally, they compared the axes of elongation of lateral connections with orientation preferences of V1 columns. In 11 of the 14 injections sites, a highly elliptical distribution of lateral connections was found (*r_n_* = 0.42) as well as a close correspondence between the axis-of-elongation of lateral connections and the preferred orientation of injected V1 columns (mean difference of 11°).

We analyzed the directional selectivity and axis-of-elongation of lateral connections in our trained models in a similar manner. Details of how we adapted the analysis for our network kernels are given in the Methods Section. *r_n_* distributions for excitatory-targeting and inhibitory-targeting kernels of a trained model are shown in Fig. 5C and 5D respectively. The average *r_n_* for excitatory-targeting kernels was found to be 0.27 while for the inhibitory-targeting kernels it was substantially lower at 0.10. Across the five trained models, we found a population-average excitatory-targeting *r_n_* of 0.25 ± 0.02 and inhibitory-targeting *r_n_* of 0.10 ± 0.01 (mean ± 1 SD). Excitatory-targeting connections were substantially more directed than inhibitory-targeting ones.

These *r_n_*s were lower than those reported in [52]. Two differences in our analysis may contribute to this. First, Sincich and Blasdel were only able to include connections that were outside a radius of 200 *μm* of the injection location, while we considered all lateral connections. Second, we weighted all lateral connections by their connection strengths so that stronger connections had a greater influence on the averaging vector, while Sincich and Blasdel considered all patch vectors to have equal weight.

Orientation differences, *θ_diff_*, between neurons’ orientation preferences and axes of elongation of their lateral connections are shown in Fig. 5E for a trained model. Each marker is scaled by the kernel’s normalized index of ellipticity, so that larger markers show more anisotropic connections. Because orientation has a period of ±180°, angular differences have a potential range of ±90°. Most neurons’ axes of elongation were close to their feedforward kernel orientations (Fig. 5E; mean excitatory-targeting *θ_diff_*=29°, mean inhibitory-targeting *θ_diff_*=31°). A smaller number of neurons had axes of elongation nearly orthogonal to their preferred orientation. The results were consistent across the 5 independently trained models (population average excitatory-targeting *θ_di_ff* = 29° ± 2° and inhibitory-targeting *θ_di_ff* = 29° ± 4°). The difference in orientations between lateral connections axis-of-elongation and feedforward orientation preferences was larger than what [52] found, but the trend was similar; most lateral connections project in the same direction as the preferred direction of their associated feedforward kernel.

Excitatory lateral connections onto inhibitory neurons in our model have a net inhibitory effect on excitatory neurons in surrounding columns. In previous contour integration models with fixed connection structures [22, 37, 38], typically a similar size is used for both excitatory and inhibitory interactions. Contrastingly, our model learnt smaller and more omni-directional inhibitory-targeting kernels. Moreover, previous models aligned the orientation of lateral inhibition kernels in the orthogonal-to-the-preferred direction of feedforward kernels, consistent with [46]. In contrast, our model learnt inhibitory-targeting connections that were mostly aligned with the preferred orientations of feedforward kernels, but more omnidirectional (Fig. 5D). These kernels are consistent with observations that short-range connections in superficial layers of V1 tend to be omni-directional and largely suppressive [53]. They are also related to a recent version of the Associate Field Model [31] that includes short-range omni-directional inhibition [54].

In summary, the lateral kernels in our model were qualitatively realistic in three respects: 1) Degree of elongation; 2) alignment of elongation with neurons’ preferred orientations; 3) relatively omnidirectional short-range inhibitory interactions. Together with realistic responses discussed in previous sections, this indicates that a physiologically realistic contour integration mechanism is consistent with optimization the contour integration network for this contour detection task.

### Edge detection in natural images

Next, we explored whether brain-like contour integration can be learnt from tasks in our natural viewing environment, and whether contour integration is useful in the performance of these tasks. Despite substantial research on the mechanisms of contour integration and the phenomenon of contour pop-out, little is known about the role of contour integration in natural life and survival. Perhaps the most specific proposal to date is that contour integration may enhance detection of parts of a contour with weak local cues, such as poor contrast [22, 37]. To test this idea, we trained our network to detect edges in natural images. We used the Barcelona Images for Perceptual Edge Detection (BIPED) dataset [55] as it considers *all contours* rather than object boundaries only. This is important because our focus is on contour integration in V1, whereas object awareness relies on more abstract representations in deeper layers. The dataset contains 200 train and 50 validation (image, edge map) pairs. It was expanded to 57,600 related training images using data augmentation methods. Sample images and ground-truth labels are shown in Fig. 6A and B.

**Fig 6.**
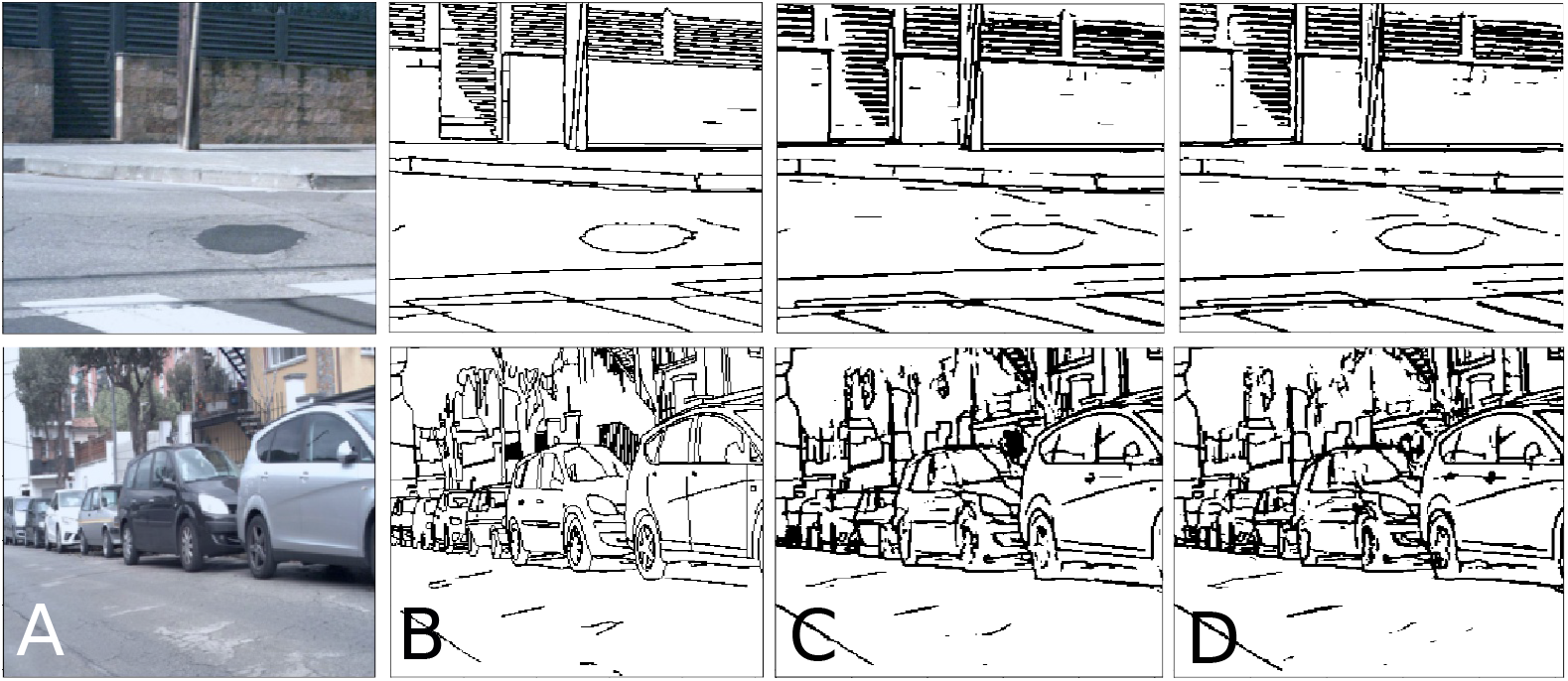
Edge detection in natural image stimuli. A, Example images from the BIPED [55] dataset. Each row shows a different image. B, Ground truth edge maps for input images shown in column A. C and D, Corresponding predictions of the control and contour-integration model, respectively.

Networks were tasked with detecting all contours in input images. Performance was evaluated using mean IoU scores (see Methods Section) between network predictions and ground-truth labels over all pixels in an image and all images in the dataset. An edge detection block (see Fig. 2) was used to map CI block outputs to the same dimensions as labels. Details of the training process are described in the Methods Section.

Example predictions of trained control and model networks are shown in Fig. 6C and D respectively. Visually, differences between their predictions are subtle. Validation IoU scores over the time course of training, for a detection threshold of 0.3 (see Methods Section), are shown in Fig. 7A. Both networks achieved their highest mean IoU scores (0.45) at this threshold. The mean IoU scores of both networks were similar. The CI block had little impact on overall performance, suggesting that the physiology of contour integration may not be essential for reliable detection of a wide variety of edges in natural scenes. To further explore this point, we trained a version of the model in which the lateral connections had a much smaller spatial extent: 3 × 3 kernels rather than 15 × 15. This model also reached the same peak performance.

**Fig 7.**
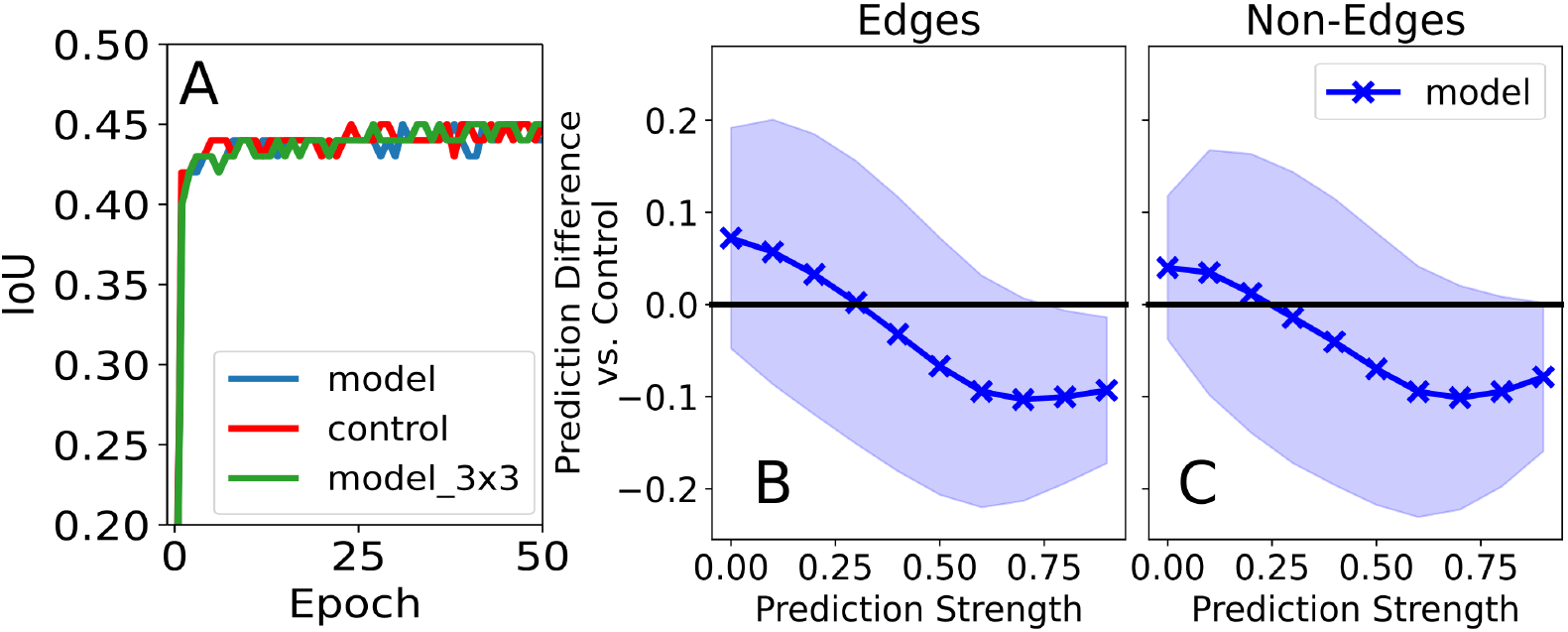
Edge detection in natural images. A, Validation IoU scores of the contour-integration model, control and a model with reduced 3 × 3 lateral kernels over training. All networks had similar performances. B and C, Average prediction difference between model and control over the validation dataset as a function of prediction strength. Model and control outputs were compared over the entire validation dataset pixel-by-pixel. Using a sliding window of width 0.2 and a step size of 0.1, control predictions within the window were highlighted and compared with corresponding model predictions. Average differences for edge pixels are shown in B and for non-edges in C. Positive values indicate higher model predictions compared to control. Solid line shows mean differences and the shaded area shows unit standard deviation around the mean.

#### Weak vs. strong edge pixel detection

In natural images, contours have non-uniform strengths and some parts are easier to detect than others. Li [37] and Piech et. al. [22] showed that contour integration can potentially enhance weak contours. However, results were only qualitatively analyzed using a single image. Although we found that contour integration did not improve detection of a wide variety of contours, including weak contours, contour integration may still strengthen low-level responses to weak contours. To investigate this question, we plotted the difference between model and control outputs as a function of the control outputs, pixel-by-pixel. Details of the procedure we used are described in the Methods Section.

The results are shown in Fig. 7B for edge pixels and in Fig. 7C for non-edge pixels. On average, the model had higher edge predictions for weaker edges (up to control output 0.3). For stronger edges, the control network responded more strongly on average. For non-edge pixels, model outputs were on average lower than control outputs for all control outputs > 0.2. This shows that model had a lower tendency toward false-positive edge detection. In summary, contour integration strengthened the representation of weak edges, but this had little practical effect on detection of weak edges at the most effective discrimination threshold.

### Naturalistic contour processing

Contour integration may support other kinds of reasoning about contours in natural scenes, for example determining which branch to climb in order to reach some fruit. To investigate this possibility, we devised a new visual perception task. Specifically, we trained the model to detect whether two points in a natural scene were part of the same contour. We placed two markers in each image. In some cases the markers were connected by a single contour in the image, while in others they were placed on different contours. We additionally punctured input images with occlusion bubbles to fragment the contours. This made it difficult to rely solely on edge extraction to solve the task. Example images are shown in Fig. 8C.

**Fig 8.**
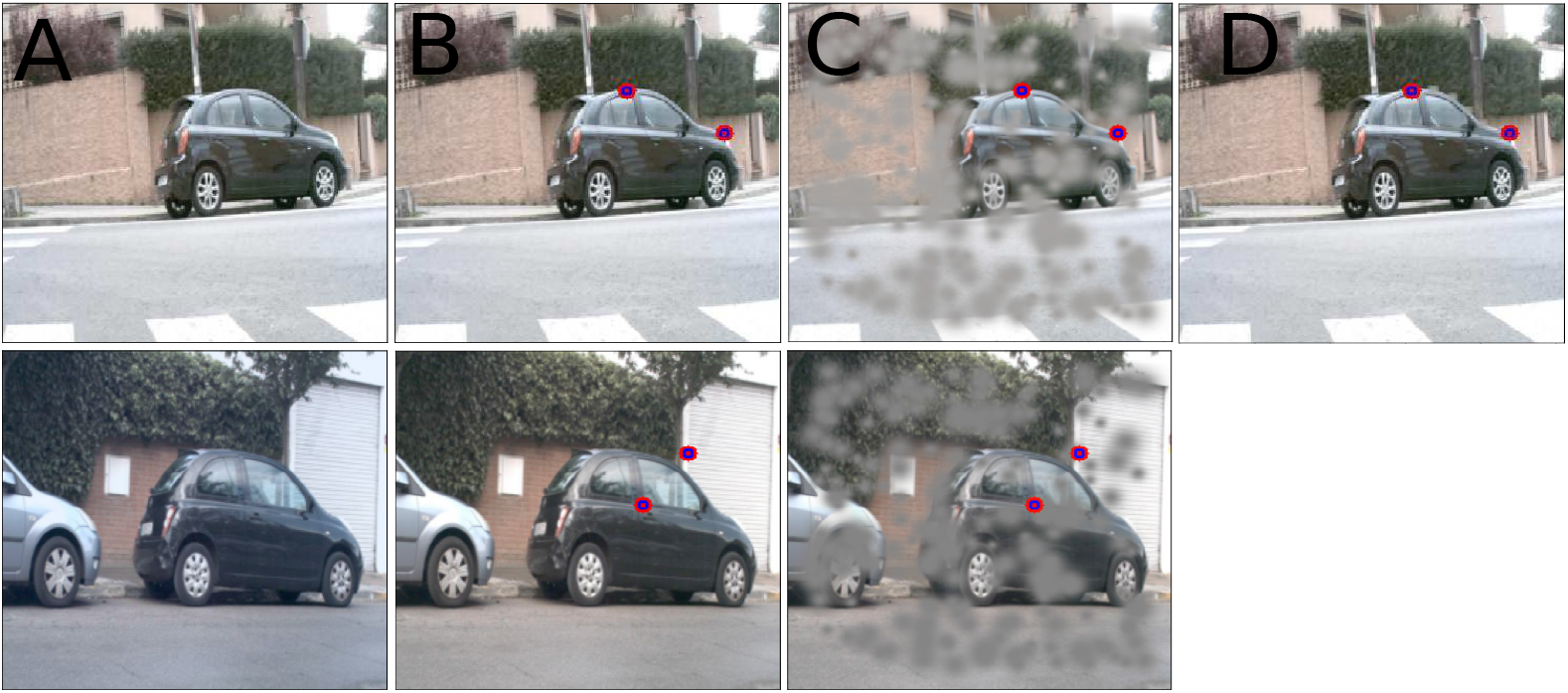
Contour tracing in natural images stimuli.. A, Sample starting image from the BIPED dataset [55]. B, Using its edges label, two markers were randomly placed on edge pixels. C, During training, images were punctured with occlusion bubbles to randomly fragment all image contours. D, After training, the impact of fragment spacing was analyzed using test contours with equidistant occlusion bubbles that were placed along contours with various inter-fragment spacing. The top row shows an example connected class stimulus, while the bottom row shows an example unconnected class stimulus.

We constructed a dataset of 50,000 training contours and 5000 validation contours that were extracted from the BIPED dataset [55]. Details of the dataset and how it was constructed are described in the Methods Section. A binary classifier block (see Fig. 2) was used to map CI block outputs to binary decisions, i.e. whether the pair of markers in each image was connected by a smooth contour or not. Performance was measured by comparing the accuracy of network predictions with labels. Training details are described in the Methods Section.

Table 2 shows peak classification accuracies averaged across five independent runs for all networks. Over the whole dataset, the model performed ≈ 5% worse than the control (validation IoU).

**Table 2.** Peak classification accuracies on the contour tracing in natural images task. Peak values (mean ± 1 SD) were averaged across five independent runs for each network. Test column shows results for test stimuli not seen during training and which had a constant inter-fragment distance of RCD = 1.

| Network | Train (%) | Validation (%) | Test (%) |
| --- | --- | --- | --- |
| Model | 70.52 ± 0.95 | 77.27 ± 1.55 | 70.39 |
| Control | 77.54 ± 0.44 | 82.67 ± 0.53 | 65.65 |

#### Effect of inter-fragment spacing

We wondered whether this natural-image task might elicit the same kinds of sensitivity to contour fragment spacing as synthetic contours [32], or whether responses to contour spacing were unique to the stimuli used in monkey experiments. We designed a variation of the task that allowed us to investigate this question in the artificial networks. To quantify the effects of inter-fragment spacing, we created new test stimuli in which inter-fragment spacing was changed in a controlled manner; occlusion bubbles were added along contours at fixed intervals. Contours were punctured with bubbles of sizes 7, 9, 11, 13, 15, 17 pixels, corresponding to fragment spacing of [7, 9, 11, 13, 15, 17]/7 RCD. An example test stimulus is shown in Fig. 8D. Details of the stimulus construction are described in the Methods Section. Binary classification accuracy was used to quantify behavioural performance while neuron responses were quantified by the contour integration gain for natural images,

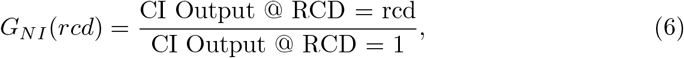

where CI Output @ RCD = 1 is the output activation of an individual neuron responding to its optimal stimulus within the cRF and with the contour fragmented with gaps the same size as the cRF stimulus, while CI Output @ RCD = rcd is the response of the neuron when the spacing between fragments was changed.

When occlusion bubbles were systematically added along contours, rather than randomly placed throughout the image, classification accuracies of all networks dropped even for the smallest bubble size (Table 2 Test column). However the relative drop in performance for the model (≈ 6%) was significantly less than that of the control (≈ 17%), showing that the strategy employed by the model generalized better from the training data to these new stimuli. Fig. 9A shows the results of fragment spacing on the behavioural performance of networks. From the least to the most spacing, model performance monotonically dropped by ≈ 4%, consistent with trends in the synthetic contour detection task. The control on the other hand, was unaffected by inter-fragment spacing.

**Fig 9.**
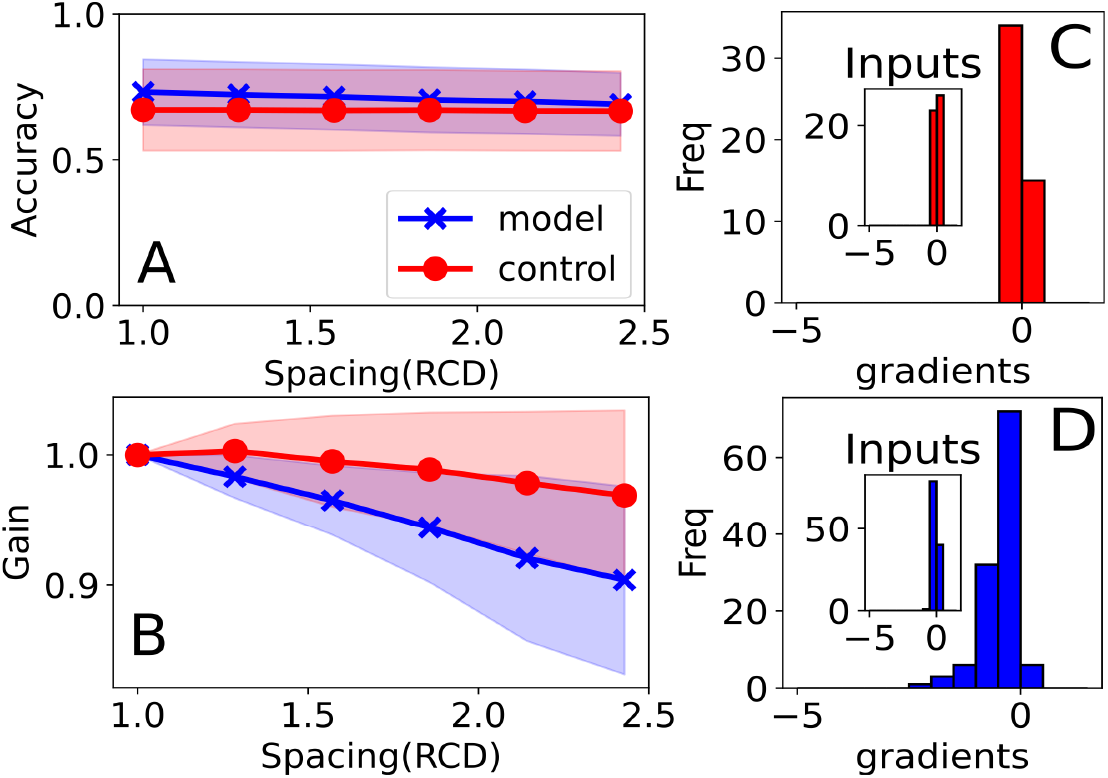
Contour tracing in natural images stimuli. A, Classification accuracy of the model (blue) and control (red) vs. fragment spacing. Dark lines show mean accuracies and shaded areas shows unit standard deviation around means. Model performance dropped as spacing increased, consistent with observed behavioural trends. B, Population average *G_NI_* vs. spacing. Gains of both networks dropped with spacing. C and D, Histograms of gradients of linear fits of gain vs. spacing results for the control and model respectively. For all networks, gains decreased with spacing. The model was more sensitive to inter-fragment spacing. Insets show histograms of gradients of CI block input activations vs. fragment spacing. Input gradients did not change significantly with spacing, showing that observed trends were learnt by CI blocks.

Fig. 9B shows population averaged contour integration gains as inter-fragment spacing increased. Population averages were found by averaging gains of individual neurons for which the optimal stimuli were found and across all five networks (trained from different random initializations) of each type. Model results were averaged across 293 neurons while control results were averaged across 120 neurons. Response gains of the control network were similar regardless of spacing, in contrast with their marked increase with spacing in the synthetic contour task. Response gains in the model decreased with increasing fragment spacing, consistent with the synthetic contour task, although the changes were less pronounced in this case.

We further analyzed the impact of fragment spacing on output activations using linear fits of output activation vs. fragment spacing of individual neurons. Histograms of the slopes are plotted in Fig. 9C and D for the control and the model networks, respectively. Similar to population averaged gain results, model outputs dropped more sharply while control output activations only dropped slightly as spacing increased.

Overall, the model behaved more consistently than the control. Its performance was less affected by new stimuli outside the training distribution, and its responses to fragment spacing were similar with both synthetic contours and natural images.

### The effect of separating excitation and inhibition

Following [22] and consistent with physiology, our model has separate excitatory and inhibitory neurons. This requires a constraint on the synaptic weights that is rarely used in deep learning and may impact performance as well as neuron responses. We created a version of the model without this constraint to test its effects. We refer to this variant as the relaxed-positivity-constraint model (RPCM). In the RPCM, each element of the lateral-connection kernels was allowed to take on any value individually. However, net lateral interactions were still restricted to be positive. This was accomplished with ReLU non-linearities operating on the weighted sums of the lateral inputs to each neuron. This is similar to the approach of [56] to model biologically plausible lateral interactions. However, individual neurons no longer conform with Dale’s principle [43, 44] as they can have both excitatory and inhibitory influences on other neurons. With this modification, the membrane potential equations of E and I nodes were defined as,

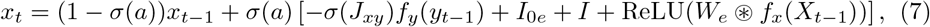

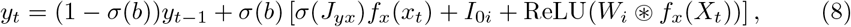

where parameters are defined in the Model Section.

On the fragmented contours dataset, the RPCM network outperformed the model by ≈ 7% and the control by ≈ 18% (Train IoU = 94.11 ± 0.05%, Validation IoU = 91.40 ± 0.12%, averaged across three networks), even though it was trained for half the time (see Methods Section). The effect of contour length on behavioural performance is shown in Fig. 10A. For all contour lengths, IoU scores of the RPCM network were higher than those of the model. Moreover, performance monotonically increased for contours of length three or longer, consistent with behavioral data. Neuron response gains also increased monotonically with contour length (Fig. 10B, results averaged over 149 neurons from three networks). However, these increases were not as pronounced as those of the model network. Similarly, RPCM neurons responded less to more widely spaced fragments, but the difference was not as pronounced as in the model network (Fig. 10C).

**Fig 10.**
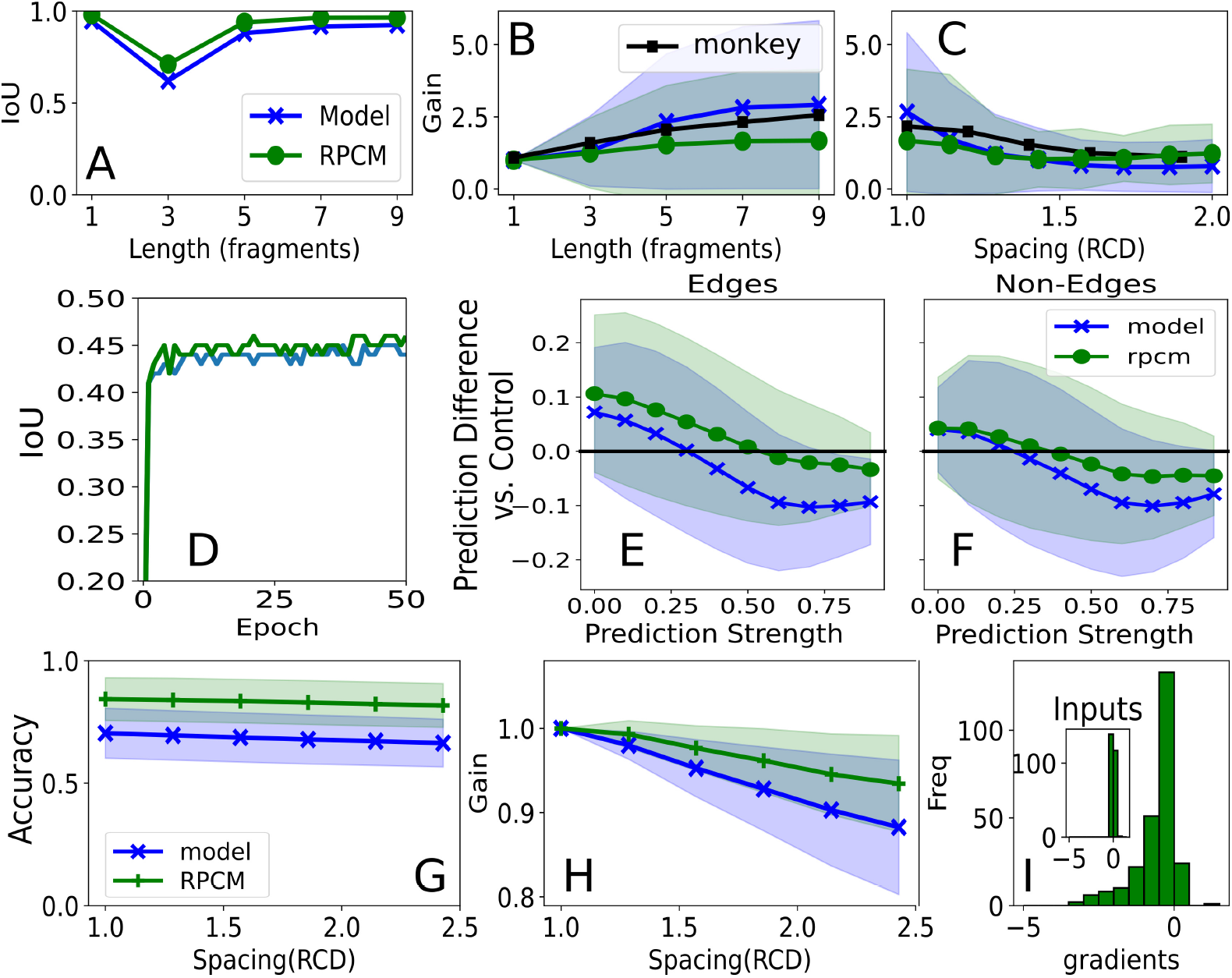
RPCM network results. The RPCM network is similar to the model but does not use a strict positive-only constraint on lateral kernels. A, IoU vs. contour length for centrally located straight contours. RPCM network IoU scores were higher than the models for all contour lengths. B and C, Population average contour integration gains vs. length and vs. fragment spacing respectively. Similar to the model, gains monotonically increased with contour length, although they were smaller. For inter-fragment spacing of up to 1.5 RCD, gains monotonically decreased. For larger spacing, gains increased slightly. D, Validation IoU scores of the RPCM network on the edge detection task in natural images. The RPCM network had a slightly higher IoU score than all other networks. E and F, Average prediction difference between RPCM and control networks for edges (E) and non-edges (F) as a function of control prediction strength. Similar to the model, the RPCM network had stronger responses to weak edges compared to the control network. (G) Classification accuracy of the RPCM network vs. fragment spacing in the contour tracing in natural images task. Performance dropped monotonically as inter-fragment spacing increased. (H) Population average vs. spacing. Neuronal gains dropped more sharply than the control but not as much as model gains.(I) Histograms of gradients of linear fits of gain vs. spacing results for the RPCM network. Similar to the model, a range of primarily negative contour integration gains were observed at the output of the CI block.

On the task of edge detection in natural images, RPCM networks peaked at a mean IoU score of 0.46 and slightly outperformed other networks (see Fig. 10D). Like contour-integration model neurons, RPCM neurons had larger responses to weak edges than control neurons (see Fig. 10E). Relative to control responses, RPCM responses varied in much the same way as model responses, although the variations were somewhat less pronounced. Similar to the contour-integration model, RPCM networks enhanced weaker contours, but this did not substantially affect task performance.

There were larger differences between the model and the RPCM on the task of contour tracing in natural images. The RPCM network outperformed the model by ≈ 13% and the control by ≈ 8% (Train 92.90 = ± 0.14, Validation = 90.62 ± 0.21). When tested with contours that were fragmented with fixed inter-fragment spacing, RPCM network performance dropped by ≈ 6% (Test = 84.36). The drop was similar to what was observed for the model and was substantially less than that of the control. RPCM networks retained the generalization properties of the model while improving overall performance. Performance also dropped monotonically with inter-fragment spacing (see Fig.10G), similar to the model. Neural response gains in the RPCM also decreased with increasing fragment spacing (Fig.10H, averaged across 257 neurons), intermediate between the model and control gains.

In summary, RPCM networks trained more quickly than contour-integration model networks and outperformed both the model and the control on every task. RPCM neurons’ responses to contour length and fragment spacing were intermediate to those of control and model neurons, but qualitatively consistent with monkey data (i.e., stronger responses with longer contours and tighter fragment spacing). Thus, Dale’s principle may have helped to account for monkeys’ neural responses, while at the same time it was functionally counter-productive in these networks and tasks.

Despite the separation of excitation and inhibition in the brain, the functional connection from any neuron to another could, in principle, be either excitatory or inhibitory depending on the strengths of direct connections and indirect connections through inhibitory interneurons [57]. We wondered whether there was a similar equivalence in our model network. Analysis of the dynamic equations (see Appendix) indicated that the contour-integration model could become functionally equivalent to the RPCM at steady state. This suggests that functional differences may be due to transient responses and/or the model being more difficult to optimize with standard algorithms in deep learning.

## Discussion

As a category, deep networks are the most realistic models of the brain, in terms of neural representations [5] and behaviour, including near-human performance on a wide range of vision tasks. However, they lack many well-known mechanisms that seem to prominently affect the function of real brains. Local circuit models [18, 19, 22, 37, 40] have the opposite limitation. They reflect specific physiological phenomena faithfully, but lack sophisticated perceptual abilities. Each of these approaches has limitations that might be alleviated by integration with the other, but such integration is rare.

Contour integration in particular has been studied extensively, but the scope of its role in visual perception is uncertain. Contour integration in V1 may occur too late [32] to drive core object recognition, which involves selectivity in inferotemporal cortex 100ms after stimulus onset [58]. It is not necessary for visual motion perception, which proceeds robustly in the absence of contours [59]. Contour integration may play a role in later stages of object recognition, together with dynamics in higher areas of the ventral stream. It could also bias core object recognition, if inferotemporal neurons learn to predict their future inputs. Such a mechanism might help to account for humans’ greater reliance on contours in object recognition compared with deep networks [9, 60]. Contour integration has been proposed to strengthen the representation of weak edges in complex scenes [22, 37]. It seems also to play a role in perceptual grouping, related to the Gestalt law of good continuation [61]. It may also be involved in segmentation, or in other kinds of reasoning about visual scenes. Integrating local circuit models into a deep networks may help to clarify the plausibility of various potential roles of contour integration in higher-level visual tasks, and may lead to new questions and predictions.

### Main findings

Our integration of a contour integration model with a deep network has produced new insights, discussed below.

#### Realistic physiology emerges from training the model to detect contours in a background of randomly oriented line segments

In contrast with past work, our model was initialized with random synaptic weights, and optimized as a whole to perform various tasks. When we trained the model to perform a contour detection task, which was similar to tasks that have been used to study contour integration in monkeys and humans, the model learned a physiologically realistic local circuit organization. Specifically, neurons in the trained model had local edge responses that were enhanced in the presence of contours, and this enhancement varied with contour length and contour fragment spacing in physiologically realistic ways. Neurons in a similar feedforward network that was trained to perform the same task did not have physiologically realistic contour responses. Their responses did not depend appreciably on contour length, and they increased instead of decreasing with contour fragment spacing. Furthermore, our contour integration model learned excitatory lateral connections that were elongated and largely aligned with neurons’ preferred orientations, as observed in the brain [52]. Past models have already established that such lateral connection patterns can produce realistic contour responses by imposing these connection patterns on the model. Our work reinforces this link by showing that it emerges consistently from an optimization process. In other words, we showed that both the lateral connections and physiological responses associated with contour integration are optimal for detecting contours in these synthetic stimuli, among a fairly generic family of networks with broad lateral connections and separate excitatory and inhibitory neurons.

#### Contour fragment spacing affects response gains similarly in natural and synthetic images

We occluded contours in natural images to test how spacing of visible contour fragments would affect contour gains. We found that greater fragment spacing monotonically reduced response strength. This result was qualitatively similar to the effect of contour spacing in synthetic images, although it was less pronounced. We do not believe that the effect of contour fragment spacing in natural images has been tested in monkeys. This would be informative, because the response patterns observed so far may only occur in response to specialized synthetic images, which would limit their ethological relevance. However, our computational results suggest that the phenomenon can generalize beyond synthetic images.

#### A contour integration model strengthens representation of edges with weak local cues in natural images

We trained the contour-integration network to detect edges in natural scenes. Compared with a feedforward control network, the contour-integration network responded more uniformly to local edge cues, with stronger responses to weak edges and weaker responses with strong edges. This confirms a suggestion by [22] and [37] that was previously only tested with a single image. However, despite these changes in local edge representation, we did not find that the contour integration model facilitated edge detection overall. The weakest edges were strengthened the most, but not enough that they exceeded the detection threshold. Indeed, because the transition from strengthening to weakening occurred near the detection threshold, and because the differences were not sufficiently pronounced (specifically, the slope in Fig. 7B was > −1), the differences in representation had little effect on edge detection. These results elaborate a previous proposal about the role of contour integration in natural images. However, while the use of natural images goes part of the way toward confronting the role of contour integration in natural life, edge detection *per se* has limited survival value. It may be fruitful in the future to consider edge representations in service of a higher-level perceptual task. In such a context, effects of contour integration below the edge detection threshold may become more relevant.

#### The contour integration mechanism can impair contour following

When we trained the model to determine whether two points in a natural image belonged to the same contour, the model performed substantially worse than the feedforward control (≈ 77% vs. ≈ 83% correct; chance performance 50%). This outcome was consistent with the impressive performance of standard convolutional networks in a wide range of vision tasks. However, it was unexpected, because the task directly involved contours. This outcome was also complicated by two factors. First, the model was better able to generalize to new stimuli than the control network. Second, the RPCM variation of the model, which did not respect Dale’s principle, outperformed the control (≈ 91% correct). The RPCM appropriately constrains the signs of net lateral influences, and exhibits physiological responses that are more realistic than those of the control network. These results indicate that recurrence in general facilitates this task, and more specifically that recurrence with some physiological properties can be beneficial. Results with the model network also show that contour integration can produce a solution that generalizes well outside the range of prior experience. However, the results do not support our expectation that physiologically realistic contour integration would improve performance of this task.

#### Dale’s principle consistently impaired performance

As a general rule, neurons release the same small-molecule neurotransmitter at each synapse (Dale’s principle), leading to distinct groups of excitatory, inhibitory, and modulatory neurons. Accordingly, our model had separate groups of excitatory and inhibitory neurons. We also tested a variant of the model (the relaxed positivity constraint model, or RPCM) that did not respect Dale’s principle but allowed the optimization process to make any synaptic weight either excitatory or inhibitory. In every task, the RPCM outperformed the more biologically grounded model. This is unsurprising because Dale’s principle amounts to a constraint on the model parameters. It is for this reason that Dale’s principle has not been adopted in deep learning.

It is unclear why Dale’s principle has been adopted in the brain, for that matter. Exceptions suggest that it could have been otherwise. For example, glutamate is normally excitatory but has inhibitory effects associated with certain receptors [62]. Some neurons elicit a biphasic inhibitory-excitatory response due to cotransmission of dopamine and GABA [63] or glutamate and GABA [64], and others change from excitatory to inhibitory depending on the presence of brain-derived neurotrophic factor [65]. So the fact that excitation and inhibition are largely separate in the brain seems to suggest that this separation is consistent with effective information processing in ways that have yet to be exploited in deep networks.

The fact that Dale’s principle impaired our model could indicate that it impairs performance of contour-related tasks in the brain, or that our model is missing other factors (e.g. feedback from higher areas, or a different kind of plasticity) that keep it from impairing performance in the brain. Consistent with the former possibility, the model that respected Dale’s principle produced the most physiologically realistic responses. However, there may be another solution that has both realistic physiology and superior task performance. Analysis of the dynamic equations indicates that the model and RPCM can become equivalent in certain conditions. This may suggest that the constrained and unconstrained models could learn similar behavior given suitable learning rules. Related to this, recent work [66] has shown that carefully designed feedforward networks with separate layers of excitatory projection neurons and intermediate inhibitory neurons can learn as well as standard deep networks, and an extension of this approach to recurrent networks was proposed. A related approach was shown to introduce new modes of instability in recurrent networks [67], but this is a promising direction for future work. Alternatively, while our model learned task-optimized lateral connections, unsupervised learning of lateral connections, as in [29], might be more effective.

### Related work

Apart from an earlier version of this work [68], our model is most closely related to the horizontal gated recurrent unit (hGRU) model [28] which similarly embeds a learnable circuit model of a low-level neural phenomenon into a larger ANN. Here we discuss some of the distinctions with that work. First, the objectives were different. Whereas we sought to test a physiologically grounded circuit model within a deep network, the purpose of the hGRU model was to improve task-level performance by using lateral connections to address the inefficient detection of long-range spatial dependencies in CNNs. Many biological constraints were relaxed to achieve higher performance. Second, the two models use different embedded circuit models. The hGRU model uses the circuit model of Mely et al. [40], a model of surround modulation, while our model uses the contour integration circuit model of Piech et al. [22]. Third, recurrent interactions in the hGRU model are derived from gated recurrent unit (GRU) networks [69]. These networks are trainable and expressive, but their internal architectures are complex and difficult to map onto circuits of the brain. Fourth, because we constrained our learnt lateral connections to be positive only, a more detailed analysis and comparison of lateral kernels was possible. In particular, we were able to compare the axis-of-elongation of lateral kernels with orientation preferences.

The V1Net model of [70] also incorporates biologically inspired lateral connection into ANNs for contour grouping tasks. The model is similar to hGRU [28] but derives its recurrent interactions from convolutional long-short-term-memory (conv-LSTM) networks [71]. Consistent with the results of the hGRU model, they find that certain recurrent ANNs, especially those with biological constraints, can match or outperform a variety of feedforward networks, including those with many more parameters. Moreover, on these tasks, they train more quickly and are more sample efficient.

## Conclusion

Local circuits are of much interest in neuroscience, but their roles in perception and behavior are mediated by the rest of the brain. Ideas about these relationships can be tested for plausibility by integrating biologically grounded models of local circuits into functionally sophisticated deep networks. Overall, our work to integrate a contour integration model into a deep network has not supported a role for this circuit in the natural-image tasks we investigated (a contour following task and detection of edges in complex natural images). This may be due to limitations of the model, although the model’s physiologically realistic responses suggest that it has much in common with the brain circuit. More work is needed to determine whether incorporating other physiological factors might produce a model that is more effective (similar to our model variant without constraints on the weight signs) without being less realistic, and to test the role of contour integration in a wider range of tasks. This line of work may be important for understanding the role of contour integration in natural life.

## Methods

### Contour integration block parameters

The architecture of the model’s contour integration (CI) block is shown in Fig. 2. In the brain, V1 lateral connections of orientation columns are sparse and preferentially connect with other orientation columns with similar selectivities [53]. Furthermore, these connections are long and can extend up to eight times the classical receptive fields (cRF) of V1 neurons [41]. Rather than using hard-coded lateral connections, we connected all columns within a *S × S* neighborhood and used task-level optimization to learn them. For edge extraction, we used the first convolutional layer of a ResNet50 [47]. It uses 7 × 7 kernels and we defined *S* to be 15 × 15. Additionally, a sparsity constraint was used during training to retain only the most important connections (see Training subsection).

Incoming feedforward signals iterated through the CI block for *N_iters_* steps before E node outputs were passed to deeper layers. We used *N_iters_* = 5 which we found to be a good trade-off between performance and run-time. Connection strengths *σ*(*J_xy_*), *σ*(*J_yx_*) were initialized to 0.1 while time constants *σ*(*a*), *σ*(*b*) were initialized to 0.5. Each neuron incorporated a rectified linear unit (ReLU) activation function, except where noted.

### Classification blocks of the network

For the task of detecting fragmented contours, CI block outputs were fed into the fragments classifier block (see Fig. 2). It consisted of 2 convolutional layers. The first convolutional layer contained 16 kernels of size of 3 × 3 and used a stride length of 1, while the second convolutional layer used a single kernel of size 1 × 1. There was a batch normalization layer between the two convolutional layers. The final convolutional layer used a logistic sigmoid non-linearity to generate prediction maps.

For the test of edge detection in natural images, CI block outputs were passed to an edge detection block (see Fig. 2). CI block outputs were upsampled by a factor of 4, using bi-linear interpolation, to resize them back to input sizes. Up-sampled activations were passed through two convolutional layers before prediction maps were generated. The first convolutional layer contained 8 kernels of size of 3 × 3 and used a stride length of 1. There was a batch normalization layer after the first convolutional layer. The last convolutional layer contained a single kernel of size 1 × 1, and was used to flatten activations to a single channel. Outputs of the final convolutional layer were passed though a logistic sigmoid non-linearity to generate prediction maps.

For the task of detecting whether two markers were connected by a smooth contour, CI block outputs were passed to the binary classifier block (see Fig. 2) that also consisted of 2 convolutional layers. The first convolutional layer consisted of 8 kernels of size 3×3 and used a stride of 3. As in the other detection blocks, there was a batch normalization layer after the first convolutional layer. The final convolutional layer used a single kernel of size 1 × 1 and used a stride of 1. Finally, a global average pooling layer [72] mapped output activations to a single value that could be compared with image labels.

### Training

Networks were trained to minimize binary cross entropy loss,

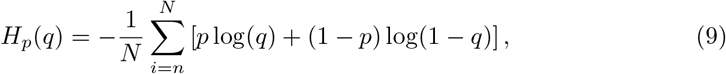

where *p* ∈ {0, 1} is the label and *q* ∈ [0, 1] is the network prediction. Here, N represents the total across all images as well as the total predictions per image.

To encourage sparse lateral connections, L1 regularization loss multiplied with an inverted 2D Gaussian mask was applied over excitatory and inhibitory lateral kernels,

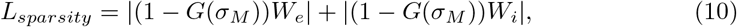

where *G*(.) is a normalized 2D Gaussian mask whose spatial spread, *σ_M_*, is defined by its standard deviation. The use of the Gaussian mask encouraged a more gradual reduction of connection strength with distance.

The total loss was defined as,

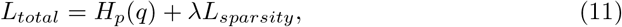

where *λ* is a weighting term for sparsity loss.

For the sparsity constraint, *σ_M_* was set to 10 pixels while *λ* was set to 1e-4. Learnt lateral connections of the model (but not the control) were restricted to be positive-only. After every weight update step, negative weights were clipped to 0.

All networks were trained with the Adam [73] optimizer. In the synthetic contour fragments detection and the contour tracing in natural images tasks, both the model and control were trained for 100 epochs with a starting learning rate (*l_r_*) of 1e-4 which was reduced by a factor of 2 after 80 epochs. The RPCM network was trained for 50 epochs with the same starting *l_r_* which was dropped by a factor of 2 after 40 epochs. Trained RPCM networks had fully converged after 50 epochs and did not noticeably improve with additional training. For edge detection in natural images, networks were trained for 50 epochs with a initial *l_r_* of 1e-3 which was reduced by a factor of 2 after 40 epochs. A fixed batch size of 32 images was used in all tasks.

All input images were fixed to a size of 256× 256 pixels, resizing images and labels when necessary. Input pixels were preprocessed to be approximately zero-centered with a standard deviation of one on average. Synthetic contour fragment images were normalized with dataset channel means and standard deviations while natural images were normalized with ImageNet values. In the contour tracing in natural images tasks, input images were punctured with occlusion bubbles as described in the Contour tracing in natural images stimuli subsection.

### Metrics

#### Mean Intersection-over-Union

For tasks with multiple outputs per image, behavioral performance was measured using the mean Intersection-over-Union metric,

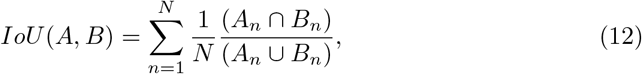

where, *N* is the number of images in the dataset, and *A* and *B* are the per-tile/pixel binary network predictions and labels for image *n*, respectively.

To get binary network predictions for an image, network outputs were passed through a sigmoid non-linearity and thresholded. The intersection with the labels was found by multiplying the predictions with their corresponding labels while the union was found by summing labels and predictions followed by subtracting the intersection of the two. An IoU score of 1 signifies a perfect match between predictions and labels, while an IoU score of 0 means that there is no match between what the network predicted and the label. Mean IoU score was found by averaging IoU scores over all the dataset.

For the contour fragments dataset a threshold of 0.5 was used, while for the contour detection in natural images tasks, a value of 0.3 returned the best scores. IoU scores dropped off monotonically as detection threshold deviated away from 0.3 for all networks.

#### Direction selectivity and axis-of-elongation of lateral connections

We followed the method of Sincich and Blasdel [52] to quantify the directional selectivity and find the axis-of-elongation of lateral connections. First, Sincich and Blasdel identified locations where stained lateral connections terminated in clusters (patches). Next, they constructed vectors originating at injection site and ending at patch centers. Given a set of patch vectors of a V1 orientation column, an averaging vector 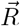 was computed. Because patch vectors pointing in opposite direction represent lateral connection extending in the same direction, the orientations of individual patch vectors were doubled before computing the vector sum. Consequently, patches that were in opposite directions summed constructively while those that were orthogonal summed destructively. After computing the vector sum, the resultant angle was halved to get the direction of the averaging vector, *θ*. To quantify directional selectivity, the magnitude of the averaging vector was normalized by the magnitude sum of all patch vectors to get a normalized index of ellipticity *r_n_*.

### Synthetic contour fragments stimuli

We used stimuli similar to those of Field et al. [31]. Each input stimulus consisted of a 2D grid of tiles that contained Gabor fragments which were identical except for their orientations and positions. The orientations and locations of a few adjacent fragments were aligned to form a smooth contour. The remaining (background) fragments had randomly varying orientations and positions.

To construct each stimulus, first, a Gabor fragment, contour length in number of fragments, *l_c_* and contour curvature, *β*, were selected. Each Gabor fragment was a square tile the same size as the cRF (kernel spatial size) of the proceeding edge extracting layer. Second, a blank image was initialized with the mean pixel value of all boundary pixels of the selected Gabor. Third, the input image was sectioned into a grid of squares (full tiles) whose length was set to the pixel length of a fragment plus the desired inter-fragment spacing, *d_full_*. The grid was aligned to coincide the center of the image with the center of the middle full tile. Fourth, a starting contour fragment was randomly placed in the image. Fifth, the location of the next contour fragment was found by projecting a vector of length *d_full_* ± *d_full_*/8 and orientation equal to the previous fragment’s orientation ±*β*. The random direction change of *β* and distance jitter were added to prevent them from appearing as cues to the network. Sixth, a fragment rotated by *β*, was added at this position. The fifth and sixth steps were repeated until ⌊*l_c_*/2⌋ contour fragments were added to both ends of the starting fragment. Seventh, background fragments were added to all unoccupied full tiles. Background fragments were randomly rotated and positioned inside the larger full tiles. Lastly, a binary label was created for each full tile indicating whether it contained the center of a contour fragment.

In all training images, inter-fragment spacing and fragment length were equal. A fixed input image size of 256×256 pixels was used. Gabor fragments of size 7×7 pixels and full tile of size 14×14 pixels were used in stimulus construction. This resulted in labels of size 19×19 for each input stimulus.

The full dataset contained 64,000 training and 6,400 validation images. In its construction, 64 different Gabors types, *l_c_* of 1, 3, 5, 7, 9 fragments and inter-fragment rotations β of 0° ± 15° were used. Gabor parameters were manually picked with the only restriction that the Gabor fragment visually appear as a well-defined line segment. Each Gabor fragment was defined over three channels and the dataset included colored as well as standard black and white stimuli. *l_c_* = 1 stimuli were included to teach the model to not do contour integration when there are no co-aligned fragments outside the cRF. Contour integration requires inputs from outside the cRF and the model had to learn when not to apply enhancement gains. For these stimuli, the label was set to all zeros. An equal number of images were generated for each condition. Due to the random distance jitter, inter-fragment rotations, and the location of contours, multiple unique contours were possible for each condition. Moreover, background fragments varied in each image.

### Test synthetic contour fragments stimuli

We used test stimuli similar to those of Li et al. [32]. These consisted of centrally located contours of different lengths and inter-fragment spacing. Test stimuli were similar to training stimuli except that the starting contour fragment was always centered at the image center. This ensured that centrally located neurons (whose outputs were measures) always received a full stimulus within their cRF. Furthermore, test stimuli were constructed in an online manner whereby the optimal stimulus of each centrally located neuron in each channel was first found by checking which of the 64 Gabor fragments elicited the maximum response in the cRF. Next, contours were extended in the direction of the preferred orientations of selected Gabors. The effects of contour length were analyzed using *l_c_* = 1, 3, 5, 7, 9 fragments and a fixed spacing of RCD=1 (see Fig. 3C). The effects of inter-fragment spacing were analyzed using RCD = [7, 8, 9, 10, 11, 12, 13, 14] / 7 and a fixed l_c_ = 7 fragments (see Fig. 3D). For each condition, results were averaged across 100 different images.

### Lateral kernel analysis

To find the direction selectivity and axis-of-elongation of lateral kernels of the model, first we found the preferred orientation of source edge extraction neurons. For each kernel in the edge extraction layer, we least-square fit each channel to a 2D Gabor function that was defined by eight parameters: the x and y location of its center, its amplitude, the orientation, wavelength and phase offset of its sinusoid component and the spatial extent and ratio of the spread in the *x* versus *y* direction of its Gaussian envelope. The orientation of the channel with the highest amplitude was selected as the kernel’s preferred orientation. Orientation preferences of the pre-trained edge extraction kernels are shown in S1 Fig. We found Gabor fits for 42 out of the 64 kernels of the edge extracting layer.

Next, following the analysis of Sincich and Blasdel [52], the average *r_n_* for each lateral kernel of the CI block was found. Excitatory-targeting (S2 Fig) and inhibitory-targeting (S3 Fig) lateral kernels were analyzed separately. Slightly different from the method of [52], individual patch vectors were calculated for every lateral weight. Moreover, as the weights of each connection were available, we weighted individual patch vectors with their connection strengths. Stronger weights contributed more to the average vector compared to weaker ones. Only those lateral kernels for which the orientation of feedforward kernels were found were considered in the analysis.

### Network predictions strengths comparison in natural images

To compare predictions of the model and the control at different edge strengths, first pre-threshold control and model outputs over the entire BIPED validation dataset were collected. Second, a sliding window of size 0.2 was run over control outputs to highlight pixels whose predictions lay within the desired range. Third, corresponding predictions of the model were found. Fourth, the average difference between model and control predictions were calculated. The process was repeated over the full range of predictions (0, 1) by sliding the window at intervals of 0.1.

Edge pixels and non-edge pixels were separately analyzed. To extract edge predictions, network outputs were multiplied with the ground truth mask. While to separate non-edge pixels, network outputs were multiplied with the inverted ground truth mask. Considering edge pixels, if the mean difference is above zero, this suggests that the model is better at detecting pixels of the corresponding strength. Considering non-edge pixels, if the mean difference is below zero, then the model has lower tendency for false positives.

### Contour tracing in natural images stimuli

The construction of stimuli for the contour tracing in natural images task required selecting contours in natural images. We randomly extracted a smooth contour *C*_1_ from a BIPED [55] image using its edge map. Contours were extracted by first selecting a random starting edge pixel from the edge map. Valid starting pixels had to be part of a straight contour in their 3× 3 pixel vicinity, either vertically, horizontally or diagonally. Next, this starting contour was extended at both ends by adding contiguous edge pixels that were at most ±*π*/4 radians from the local direction of the contour. The local direction of the contour was defined as the angular difference between the last two points of the contour. If there was more than one candidate edge pixel, the candidate with the smallest offset from the contour direction was selected. The process was repeated until there were no more edge pixels at candidate positions or if the selected candidate pixel was already a part of *C*_1_ (circular contours). Additionally, once contour length was greater than 8 pixels, a large-scale smooth curvature constraint was applied to check that the angle difference between (*n, n* – 4) and (*n* – 4, *n* – 8) contour points was not greater than *π*/4 radians, where, n is the last point on the contour. Contour extraction was also stopped if the large-scale curvature constraint was not met.

After extracting *C*_1_, one of its endpoints was chosen as the position of the first marker, *M*_1_. Next, a second edge pixel that did not lie on *C*_1_ was randomly selected. To ensure that connected and unconnected stimuli had similar separation distances, the selection process used a non-uniform probability distribution to favor edge pixels that were equidistant with the unselected endpoint of *C*_1_. First, distances for all edge pixels from *M*_1_ were calculated. Next, the absolute difference between edge pixels distances and the distance to the unselected endpoint of *C*_1_ was calculated. A Softmax function was used to convert negative distances differences to probabilities. Edge pixels that were of a similar distance to the unselected end point of *C*_1_ had distance differences close to zero and were more likely to be selected, while edge pixels that were at a different distance had large negative distance differences and were less probable.

Given the location of the second edge pixel, a second contour, *C*_2_, was extended from it. If any point on *C*_2_ overlapped with *C*_1_, a new starting edge pixel was selected and the process was repeated until a non-overlapping pair of contours was found. The location of the second marker, *M*_2_, was determined by the type of stimulus. For connected stimuli, the opposite end of *C*_1_ was selected as *M*_2_, while for unconnected stimuli, one of the endpoints of *C*_2_ was chosen. Once marker positions were determined, markers were placed at corresponding positions in the input image. Each marker consisted of a bulls-eye of alternating red and blue concentric circles (see Fig. 8B). Markers were added directly to input natural images, and networks were given no information about the selected contours.

To fragment contours, occlusion bubbles were added to input images. Following [74], bubbles with a 2D Gaussian profile were used to reduce the impact of bubble edges. Each image was punctured using 200 bubbles of multiple sizes. Bubble sizes were specified by the full-width-half-maximum (FWHM) of 2D Gaussian functions and were chosen to correspond to bubble sizes used to explore the effects of fragment spacing on neurophysiological gains (see subsection Test contour tracing in natural images stimuli). Individual bubbles were defined over a 2× FWHM square area. After randomly selecting bubble sizes and locations, bubbles were placed in a mask which was used to blend the input image with image channel mean values using,

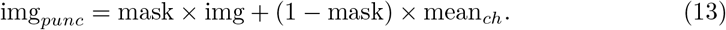

Within a mask, bubbles were allowed to overlap and a different mask was used for each image. Values in the bubble mask ranged between [0, 1]. Sample input training images for the contour tracing task are shown in Fig. 8C.

The train dataset contained 50,000 contours that were extracted from BIPED train images while the validation dataset contained 5,000 contours that were extracted from BIPED test images. Since the BIPED test dataset contains only 50 images, multiple contours per image were extracted. Care was taken to ensure duplicate contours were not selected. Puncturing of input images was done as a pre-processing step during the training loop. Consequently, each exposure of an image to a network was unique. Equal probabilities were used for generating connected and unconnected stimuli.

### Test contour tracing in natural images stimuli

Similar to when the effects of inter-fragment spacing were analyzed using synthetic fragmented contours, the optimal stimuli of target neurons needed to be found. In the synthetic contour fragments dataset, test images were designed to contain the optimal stimuli of monitored neurons. However, for natural images, inputs cannot be defined in a similar way. Therefore, a new procedure was devised. To find the optimal stimulus of an individual channel, multiple unoccluded connected contours were presented to networks (Fig. 8B). For each image, the position of the most active neuron of each channel in the CI block was found. If it was within 3 pixels (the same as the stride length of the subsequent convolutional layer) of the contour, the image as well as the position of most active neuron were stored. he process was repeated over 5,000 contours and the top 50 (contour, most active neuron) pairs were retained for each channel. New random contours were selected from the augmented BIPED train dataset. The train dataset, as opposed to the test dataset, was used as it contained more images and a larger variety of contours.

Given the optimal stimulus for a channel, each input contour was fragmented by inserting occlusion bubbles at specific positions along the contour. Different bubble sizes were used to fragment contours with different inter-fragment spacing. A fixed fragment length of seven pixels, the same size as the cRF of edge extracting neurons, was used. To ensure the cRF of the most active neuron was unaffected by bubbles, first, the position of the closest point on the contour was found. Bubbles were then inserted along the contour at offsets of ± (*l_frag_ + l_hubble_*)/2, ±3(*l_frag_ + l_bubble_*)/2, ±5(*l_frag_ + l_bubble_*)/2,… until the ends of the contour. Finally, the blending-in area of bubbles was restricted to FWHM pixels to ensure visible contour fragments were unaffected.

## Code availability

Source code for all networks, experiments and analysis that were performed as well as for generating datasets used in this work is available at https://github.com/salkhan23/contour_integration_pytorch.

## Appendix: The effect of Dale’s principle on model function

The model’s dynamic equations are,

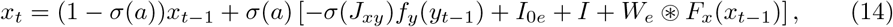

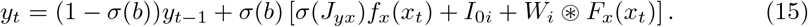

The term inside square brackets in (1) is the drive into *x*,

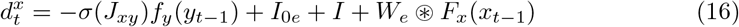

*y* does not receive inhibitory input, so if *I*_0*i*_ is positive then *y* is positive, and the rectifying function *f_y_* can be ignored. Suppose we set *σ*(*J_xy_*) = 1. Under these conditions *d* can be simplified to,

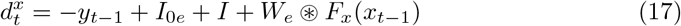

As *y_t_* approaches steady state,

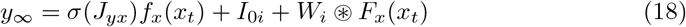

We can set *σ*(*J_yx_*) = 0 by absorbing this factor into *W_i_*. Then,

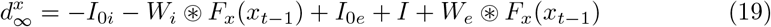

In summary, *W_i_* and *W_e_* affect *x_t_* in the same way in the following conditions: 1) when *y* reaches steady state; 2) assuming *I*_0*i*_ ≥ 0; 3) *σ*(*J_xy_*) = 1; 4) *σ*(*J_yx_*) = 0. So in these conditions, if both matrices were unconstrained and contained both positive and negative values, Dale’s Principle could be re-established by moving the positive values to *W_e_* and the negative values to *W_i_*, without change of function.

More generally, the latter two factors do not have to be enforced. If *σ*(*J_yx_*) > 0 then it can be moved into the diagonal of *W_i_*. Similarly, if *σ*(*J_xy_*) = *g* < 1 then it can be multiplied by *1/g* and *W_i_* and *I*_0*i*_ multiplied by *g*, without changing the function.

This suggests that differences between model and RPCM are due to transient dynamics and learning dynamics (i.e. the model may be structurally capable of RPCM performance but the solution may not be reachable via backpropagation and Adam).

## Supporting information

**S1 Fig.**
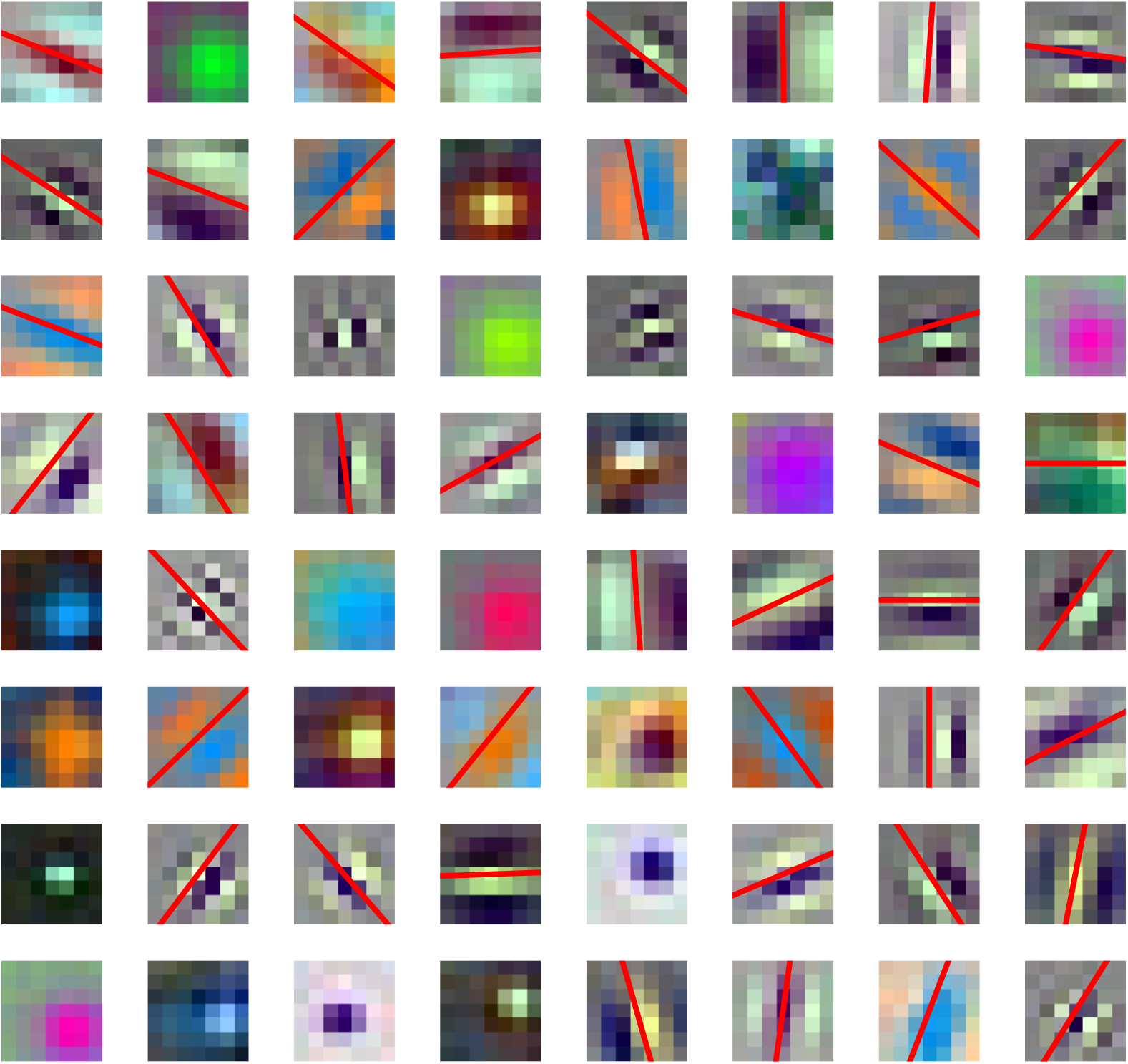
Feedforward edge extraction kernels and their preferred orientations. Each subplot shows one of the 64 kernels of the first convolutional layer of a ResNet50 model that was trained on ImageNet [48]. This served as the main component of the edge extraction block. It contains 64 kernels and each kernel has 3 input channels and a spatial spread of 7×7 pixels. Each kernel was fit to a 2D Gabor function to find its preferred orientation (red lines). The fitting algorithm, was able to find the orientation preferences of 42 kernels. Kernels for whom no fits were found (no red line) were skipped and not used in the analysis.

**S2 Fig.**
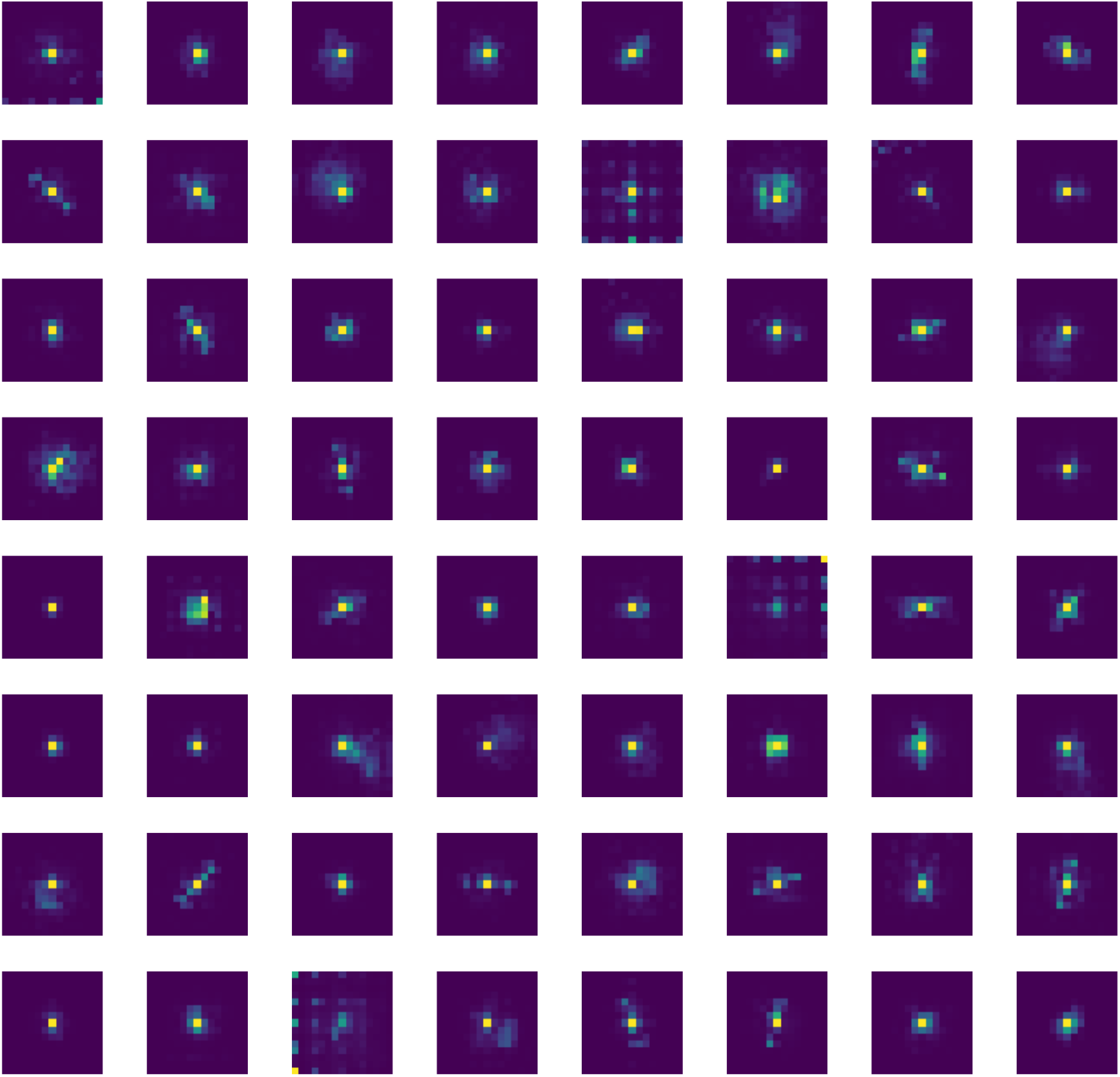
Learnt lateral excitatory kernels. Each subplot plots one of the 64 learnt excitatory lateral kernels of a model trained on the synthetic contour fragments dataset. Individual kernels had 64 channels and had a spatial spread of 15 × 15. To view the kernels, the channel dimensions were compressed by summing over all channels. Many excitatory kernels appear to be highly directed, spreading out in one dimension more than others.

**S3 Fig.**
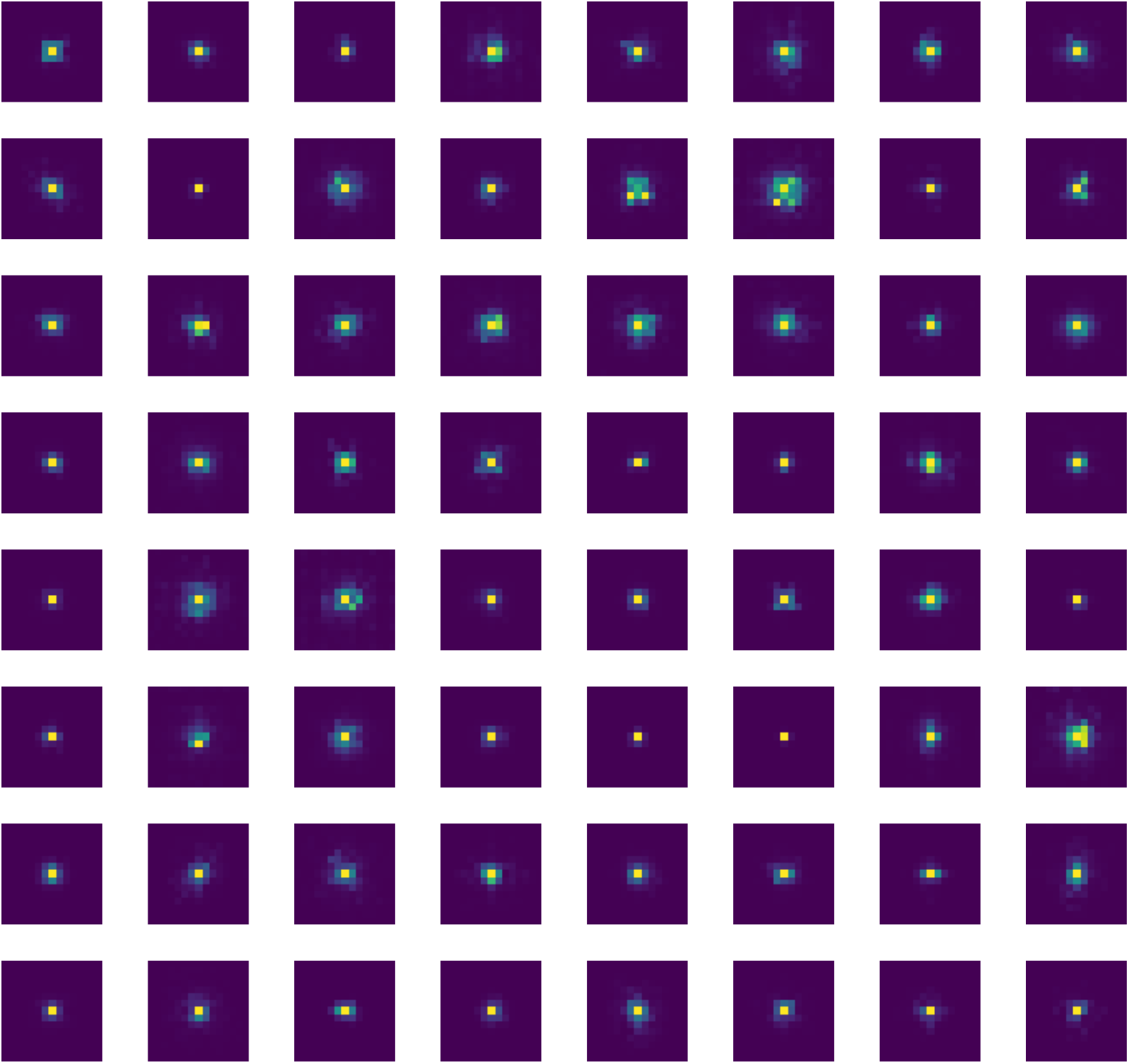
Learnt lateral inhibitory kernels. Each subplot plots one of the 64 learnt inhibitory lateral kernels of a model trained on the synthetic contour fragments dataset. Individual kernels had 64 channels and had a spatial spread of 15 × 15. To view the kernels, the channel dimensions were compressed by summing over all channels. The spatial extent of inhibitory kernels was less than the spread of excitatory kernels and mostly omni-directional.

